# Disrupted Mechanobiology Links the Molecular and Cellular Phenotypes in Familial Dilated Cardiomyopathy

**DOI:** 10.1101/555391

**Authors:** Sarah R. Clippinger, Paige E. Cloonan, Lina Greenberg, Melanie Ernst, W. Tom Stump, Michael J. Greenberg

## Abstract

Familial dilated cardiomyopathy (DCM) is a leading cause of sudden cardiac death and a major indicator for heart transplant. The disease is frequently caused by mutations of sarcomeric proteins; however, it is not well understood how these molecular mutations lead to alterations in cellular organization and contractility. To address this critical gap in our knowledge, we studied the molecular and cellular consequences of a DCM mutation in troponin-T, ΔK210. We determined the molecular mechanism of ΔK210 and used computational modeling to predict that the mutation should reduce the force per sarcomere. In mutant cardiomyocytes, we found that ΔK210 not only reduces contractility, but also causes cellular hypertrophy and impairs cardiomyocytes’ ability to adapt to changes in substrate stiffness (e.g., heart tissue fibrosis that occurs with aging and disease). These results link the molecular and cellular phenotypes and implicate alterations in mechanosensing as an important factor in the development of DCM.

## Introduction

DCM is a major cause of sudden cardiac death in young people, and it is a significant cause of heart failure (1). DCM is phenotypically characterized by dilation of the left ventricular chamber, and it is often accompanied by changes in cellular and tissue organization, including lengthening of individual myocytes and fibrosis-induced stiffening of the myocardial tissue. Genetic studies have demonstrated that familial DCM can be caused by single point mutations, many of which are in sarcomeric proteins responsible for regulating cardiac contractility (1); however, the connection between the initial insult of molecular-based changes in contractile proteins and the development of the cellular disease phenotype is not thoroughly understood. This lack of understanding has hampered efforts to develop novel therapeutics, and there is currently no cure for DCM, with heart transplantation being the only long-term treatment. The goal of this study was to better understand the connection between mutation-induced changes in contractility and the cellular phenotype.

Understanding the link between point mutations in sarcomeric proteins and the development of the disease phenotype in cells has been challenging for several reasons. One challenge stems from the fact that the clinical presentation and prognosis appear to depend on the specific mutation. In fact, different point mutations within the same molecule, and even different substitutions at the same amino acid site can lead to different forms of cardiomyopathy (e.g., hypertrophic, dilated, restrictive) with different ages of onset (2–5). Another challenge stems from selecting appropriate model systems of the human disease for the specific question being posed. *In vitro* biochemical studies are excellent for deciphering the molecular consequences of the initial insult (4); however, they are not clearly predictive of how the disease will manifest itself in cells (6). Studies using transgenic mice have significantly advanced our understanding of the disease; however, these mice do not always recapitulate the human disease phenotype due to differences in cardiac physiology between humans and mice (7–13). Patient tissue is excellent for studying the disease phenotype in humans (14); however, tissue from patients is difficult to obtain and usually only available from patients in the end stages of the disease or post mortem, when compensatory mechanisms can mask the initial insult.

Recent advances in stem cell (15, 16) and gene editing technologies (17) have provided a new avenue for modeling DCM in human pluripotent stem cell derived cardiomyocytes (hPSC-CMs) (18, 19). These hPSC-CMs can faithfully recapitulate many aspects of the early human disease (20), making them an excellent tool for deciphering how the initial molecular insult leads to the development of the early disease phenotype. Moreover, these cells can be examined in simplified *in vitro* systems that mimic various aspects of the environment in the heart to dissect how the local environment influences the development of the disease phenotype. It has been shown that cardiomyocytes can adapt their contractility in response to their local mechanical and geometric environment (21–23). It is possible aberrant responses of cardiomyocytes to disease-related alterations in the local environment such as fibrosis-induced stiffening of the heart tissue could contribute the development of the disease phenotype.

To uncover the connection between molecular changes and the development of the cellular disease phenotype, we determined the molecular and cellular mechanisms of a DCM-causing point mutation in the protein troponin-T (TNNT2), deletion of lysine 210 (ΔK210) (24) (Figure 1A). The ΔK210 mutation has been identified in patients as young as 2 years old in at least four unrelated families (25–27). Troponin-T is a subunit of the troponin complex, which, together with tropomyosin, regulates the calcium-dependent interactions between the force-generating molecular motor myosin and the thin filament. TNNT2 is one of the most frequently mutated genes in DCM (28, 29). There have been several excellent model systems developed to better understand aspects of the disease caused by this mutation (30–34). However, the connection between the molecular mutation and the cellular phenotype seen in humans remains unclear.

**Figure 1:**
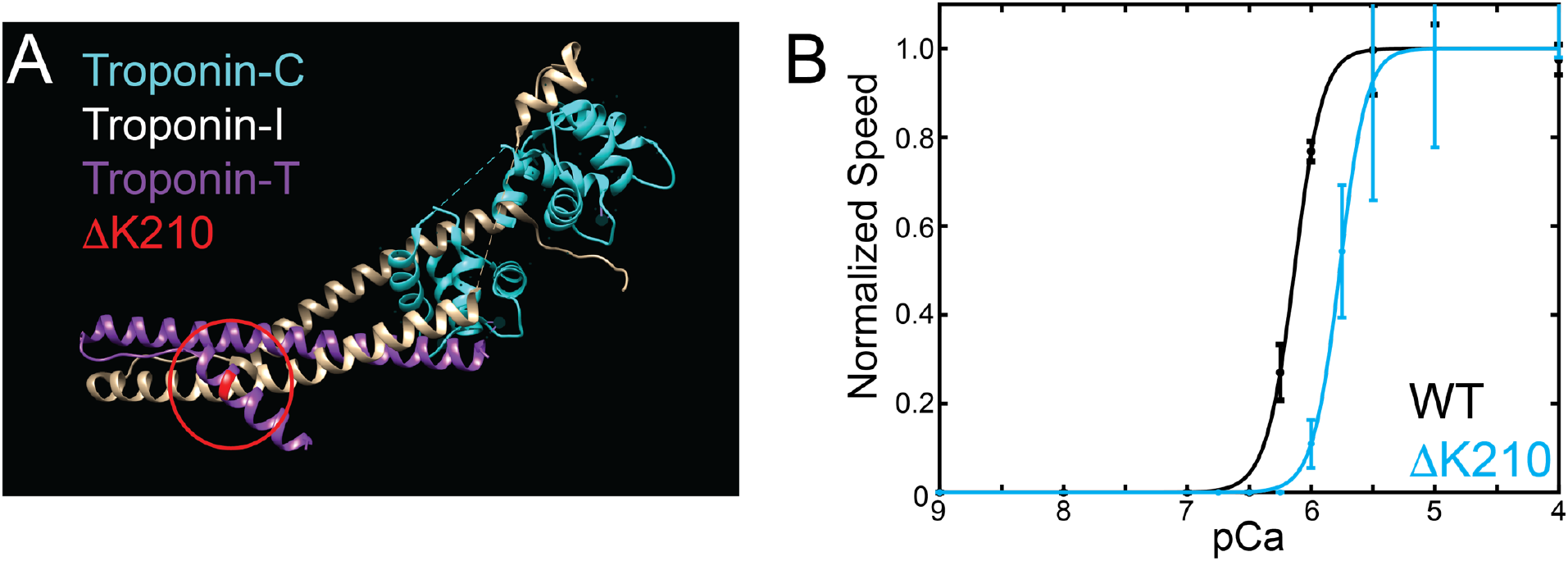
(A) Structure of the troponin core complex (PDB: 4Y99), showing troponin-C (cyan), troponin-I (white), troponin-T (purple), and K210 of troponin-T (red). (B) Speed of thin filament translocation as a function of calcium measured in the *in vitro* motility assay. Error bars show the standard deviation of n=3 experiments. Data show a significant shift in the pCa50 towards submaximal calcium activation for ΔK210 (p<0.0001). The Hill coefficient is not changed between the WT and ΔK210 (p = 0.45).

Our results demonstrate that ΔK210 causes a reduction in myosin-based force generation at the molecular and cellular levels, and we used computational modeling to link the contractile phenotypes seen at these scales. Surprisingly, we found that this disease-causing mutation of a sarcomeric protein not only impairs cardiomyocyte contraction, but it also causes cellular hypertrophy and impairs the ability of cardiomyocytes to sense and respond to changes in their mechanical environment. Our results suggest a central role for defects in mechanosensing in the disease pathogenesis.

## Results

### ΔK210 decreases calcium sensitivity in an *in vitro* motility assay

We set out to decipher the molecular mechanism of the ΔK210 mutation *in vitro.* The molecular effects of cardiomyopathy mutations depend on the myosin isoform (7–9, 35–37) and therefore, we used porcine cardiac ventricular myosin (38). Porcine ventricular cardiac myosin (MYH7) is 97% identical to human, while murine cardiac myosin (MYH6) is only 92% identical. Porcine cardiac myosin has very similar biophysical properties to human cardiac myosin, including the kinetics of the ATPase cycle, step size, and sensitivity to load (38–41), making it an ideal myosin for biophysical studies.

Given the role of troponin-T in thin filament regulation, we first determined whether the ΔK210 mutation affects calcium-based regulation of myosin binding to thin filaments using an *in vitro* motility assay (42). Reconstituted thin filaments, consisting of porcine cardiac actin and recombinantly expressed human troponin and tropomyosin, were added to a flow cell coated with porcine cardiac myosin in the presence of ATP. The speed of filament translocation was measured as a function of added calcium. As has been reported previously, the speed of regulated thin filament translocation increased sigmoidally with increasing Ca^2+^ concentration (43), (Figure 1B). Data were fit with the Hill equation to obtain the pCa50 (i.e., the concentration of calcium necessary for half-maximal activation). Consistent with previous studies using mouse cardiac, rabbit cardiac, and rabbit skeletal muscle fibers (31, 33, 44), ΔK210 shows a right-shifted curve (pCa50 = 5.7 ± 0.1) compared to the WT (pCa50 = 6.1 ± 0.1; p < 0.0001), meaning more calcium is needed for the same level of activation. This suggests that the mutant could show impaired force production during a calcium transient.

### Molecular mechanism of ΔK210-induced changes in thin filament regulation

The changes in calcium sensitivity seen in the *in* vitro motility assay could be due to alterations in thin-filament based regulation or allosteric effects on myosin’s kinetics. To test whether the mutation affects the kinetics of myosin dissociation from thin filaments, we measured the rate of ADP release from myosin, the transition that limits actomyosin dissociation at saturating ATP in the absence of load (45). Consistent with previous studies of other DCM troponin mutations (46), ΔK210 does not change the kinetics of ADP release from myosin, (Figure 1 – figure supplement 1), and therefore, the primary effects of the mutation are likely due to alterations in thin filament-based regulation.

Biochemical (47) and structural biological (48, 49) experiments have shown that thin filament regulation is a complicated process where tropomyosin can lie along the thin filament in three positions (blocked, closed, and open), and that this positioning is determined by both calcium and myosin binding (Figure 2A). To determine the molecular mechanism of the observed changes in the calcium sensitivity of myosin-based motility, we determined the equilibrium constants for the transitions between these states.

**Figure 2:**
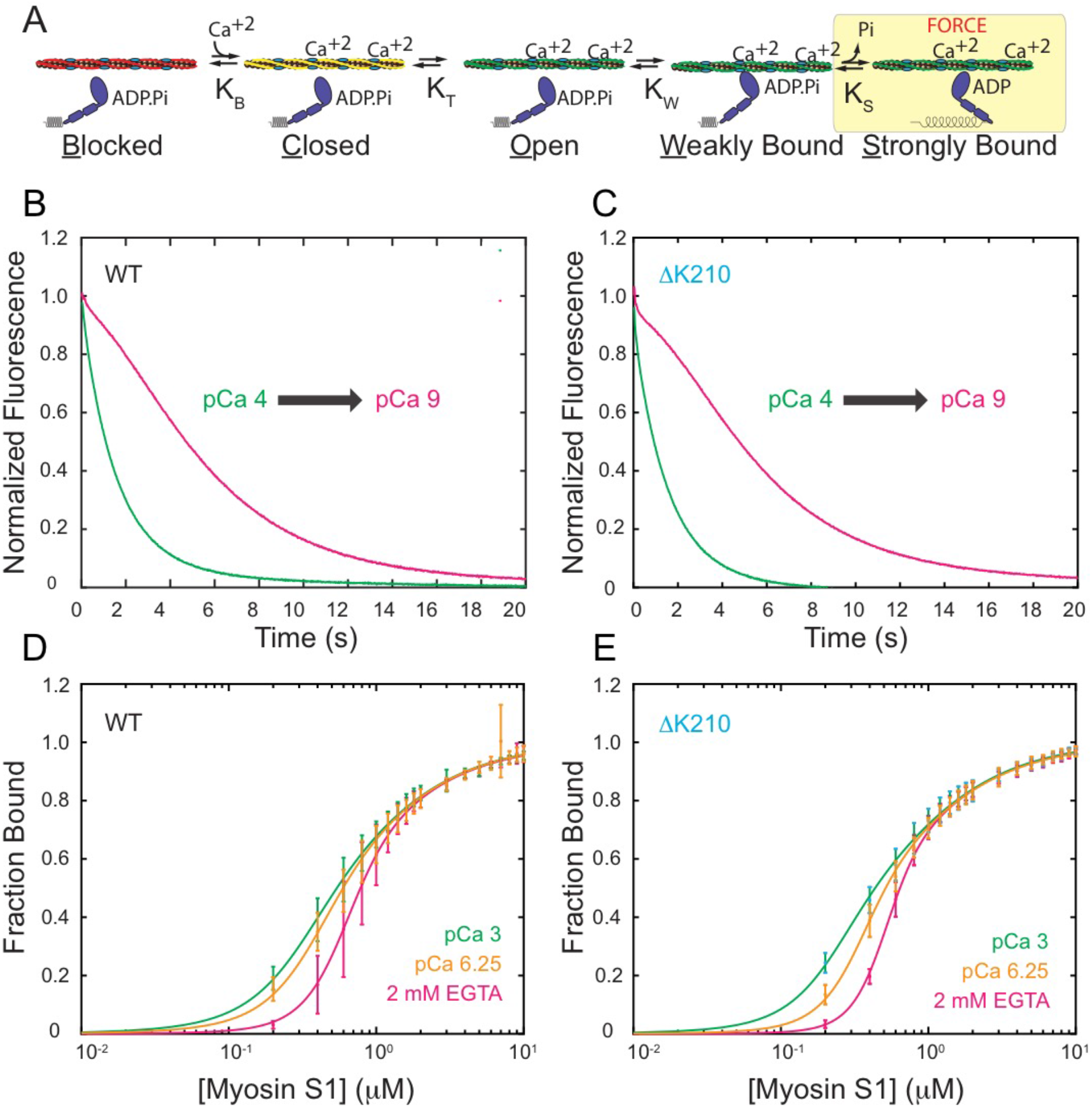
Determination of regulated thin filament parameters for ΔK210. (A) The three-state model for tropomyosin positioning along the thin filament (47). Activation of the thin filament and subsequent force generation requires both calcium and myosin binding. The equilibrium constant for the blocked to closed transition is *K*_B_ and the equilibrium constant for the closed to open transition is *K*_T_. *K*_W_ and *K*_S_ are the equilibrium constants for myosin weak- and strong-binding to the thin filament, respectively. (B-C) Normalized stopped-flow fluorescence traces of myosin binding to regulated pyrene-actin binding. The quenching of pyrene-actin upon binding of S1 occurs at a faster rate at pCa 4 (green) compared to pCa 9 (pink) due to the activating effect of calcium. Traces for (A) WT and (B) ΔK210 are very similar and the calculated values of *K*_B_ are not statistically different (p=0.998). (D-E) Fluorescence titrations in which increasing amounts of S1 myosin were gradually added to regulated thin filaments to a final concentration of 10 μM S1 in the presence of ADP. Titration curves at three different calcium concentrations are shown for both (C) WT and (D) ΔK210. 5 repeats were performed for each condition. At low concentrations of myosin, more myosin is bound to the thin filament at higher calcium concentrations (pCa 6.25 (orange) and pCa 3 (green)) than at low calcium (2 mM EGTA, pink).

The equilibrium constant for the transition between the blocked and closed states, *K*_B_, was measured using stopped-flow kinetic techniques (47), where the rate of myosin binding to pyrene-labeled regulated thin filaments was measured at high (pCa 4) and low (pCa 9) calcium concentrations. Myosin strong binding to labeled thin filaments quenches the pyrene fluorescence, leading to a roughly exponential decrease in fluorescence (Figure 2B-C). The ratio of the rate constants for myosin binding to the thin filament at high and low calcium can be used to calculate the equilibrium constant, *K*_B_ (47):

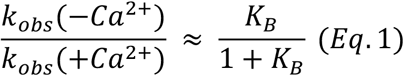

The values of K_B_ obtained were 0.36 ± 0.15 for WT (n=3) and 0.36 ± 0.10 for ΔK210 (n=3), indicating that ΔK210 does not significantly affect the equilibrium constant for the transition between the blocked and closed states (p = 0.99).

To determine the equilibrium constant for the transition between the closed and open states, *K*_T_, we measured the steady-state binding of myosin to pyrene-labeled regulated thin filaments over a range of myosin concentrations (Figure 2D-E). Titration relationships obtained for WT and ΔK210 at high (pCa 3) and low (2 mM EGTA) calcium concentrations show two-state (i.e., no calcium-based blocking) and three-state processes, respectively, as has been observed previously (47). Five technical replicates were used to define each curve. As described in the Materials and Methods, we modified the fitting and analysis procedure used by McKillop and Geeves to better define the values of the fitted variables (50). We performed an additional titration at an intermediate calcium concentration (pCa 6.25) and used an annealing algorithm to globally fit the binding curves of the fractional change in pyrene fluorescence (α) as a function of myosin concentration for all three calcium concentrations (Figure 2D-E). The fractional change in the pyrene fluorescence is given by (47):

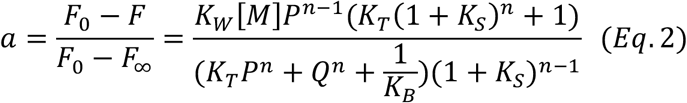

where [M] is the concentration of myosin, *F* is the measured fluorescence, *F*_0_ is the fluorescence in the absence of myosin, *F*_∞_ is the fluorescence at saturating myosin concentrations, *n* is the size of the cooperative unit, *P*=1+*K*_W_[M](1+*K*_s_), and Q=1+*K*_W_[M] (Figure 2D-E). For the fitting, *K*_T_, *K*_W_ (i.e., the equilibrium constant between the open and myosin weakly bound states, sometimes denoted *K*_1_), and *n* were fit parameters. *K*_B_ and *K*_S_ (the equilibrium constant between the myosin weakly and strongly bound states, sometimes denoted *K*_2_) were fixed based on the calculated values for *K*_B_ (Figure 2B-C) and previously measured values of *K*_S_ (51). 95% confidence intervals were calculated from 1000 rounds of bootstrapping simulations (see Materials and Methods).

We found that while the measured values of *K*_T_ for ΔK210 were consistently lower compared to the WT at all calcium concentrations (Figure 3A), only the difference in *K*_T_ at pCa 6.25 was statistically significant (p=0.028). From the measured equilibrium constants, it is straightforward to calculate the fraction of regulatory units in each state from the partition function. At all calcium concentrations, ΔK210 shows a decrease in the fraction of thin filaments in the open, weakly-bound, and strongly-bound states compared to the WT (Figure 3B), and an increase in the fraction of thin filaments in the blocked and closed states. These data give the biochemical mechanism of the reduced activation at sub-saturating calcium seen in the *in vitro* motility assays (Figure 1B).

**Figure 3:**
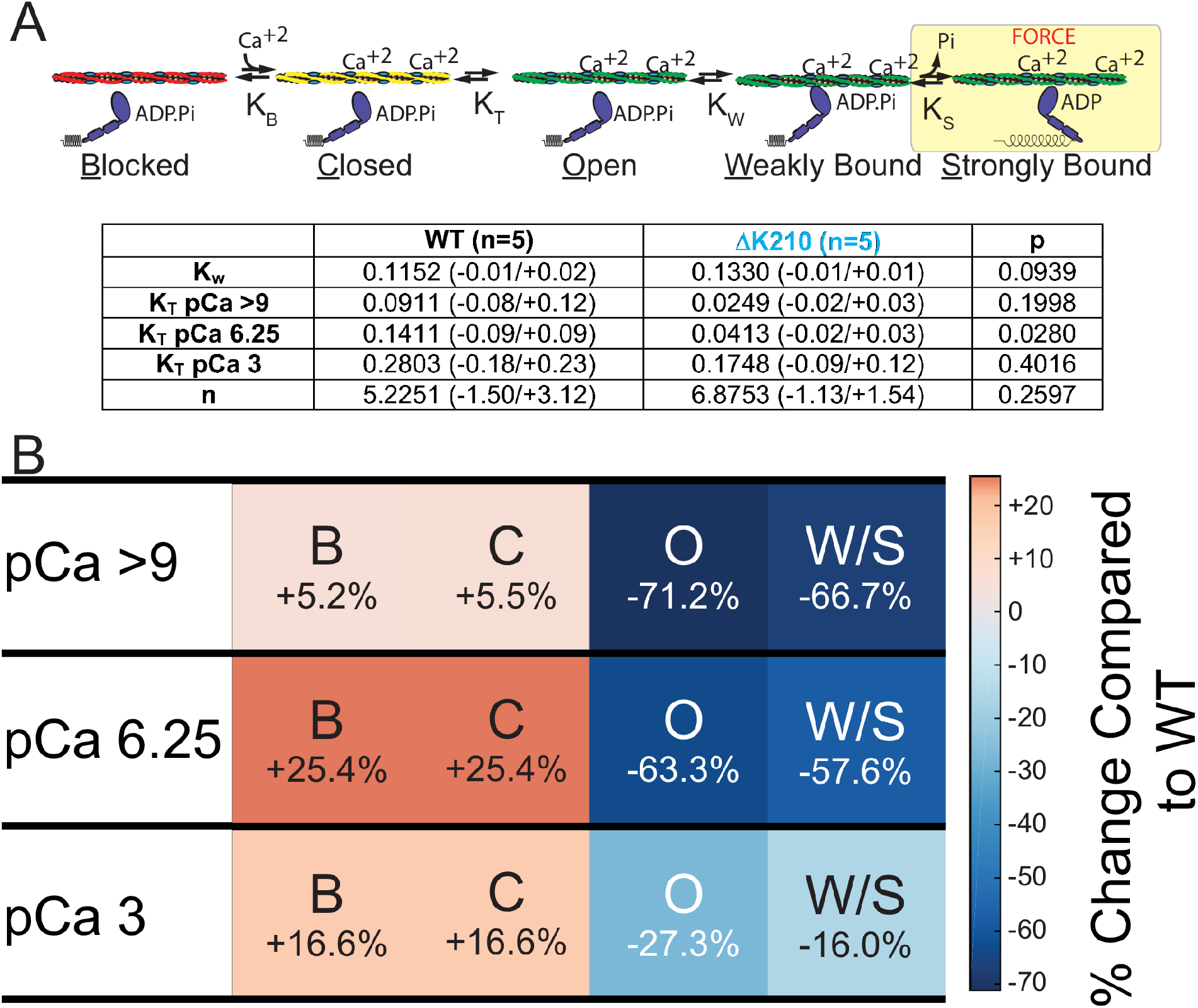
Effects of ΔK210 on tropomyosin positioning along the thin filament. (A) The three-state model for tropomyosin positioning along the thin filament (47) and parameter values obtained from the stopped-flow and fluorescence titration experiments. *K*_T_ for ΔK210 at pCa 6.25 is statistically different from WT (p=0.028). Error bars are the 95% confidence intervals. (B) Percent change in the occupancy of each state for the mutant compared to the WT. Reds denote increased population of a state, while blues denote decreased population. ΔK210 causes a decrease in the population of the states where myosin is bound to the thick filament and an increase in the population of the inhibited (i.e., blocked and closed) states. Note that the change in the fractions in the blocked and closed states is always the same, since *K*_B_ does not change between the WT and the mutant.

### Computational modeling recapitulates the shift in calcium sensitivity and predicts a lower force per sarcomere for ΔK210

To connect how the observed changes in the equilibrium populations of thin filament regulatory units with contractility in a sarcomere, we used a computational model developed by Campbell et al. (52). Given a set of equilibrium constants for thin filament activation, this model calculates both the steady-state force as a function of calcium and the force produced per sarcomere in response to a calcium transient. Using our measured equilibrium constants, the steady-state normalized force was simulated for both WT and ΔK210 over a range of calcium concentrations (Figure 4A). Simulation of ΔK210 shows a rightward shift towards supermaximal calcium activation, consistent with the shift seen in the *in vitro* motility assay (Figure 1B). Therefore, the observed set of changes in the equilibrium constants in the mutant can mechanistically explain the observed changes seen in the *in vitro* motility assay, validating the approach taken here.

**Figure 4:**
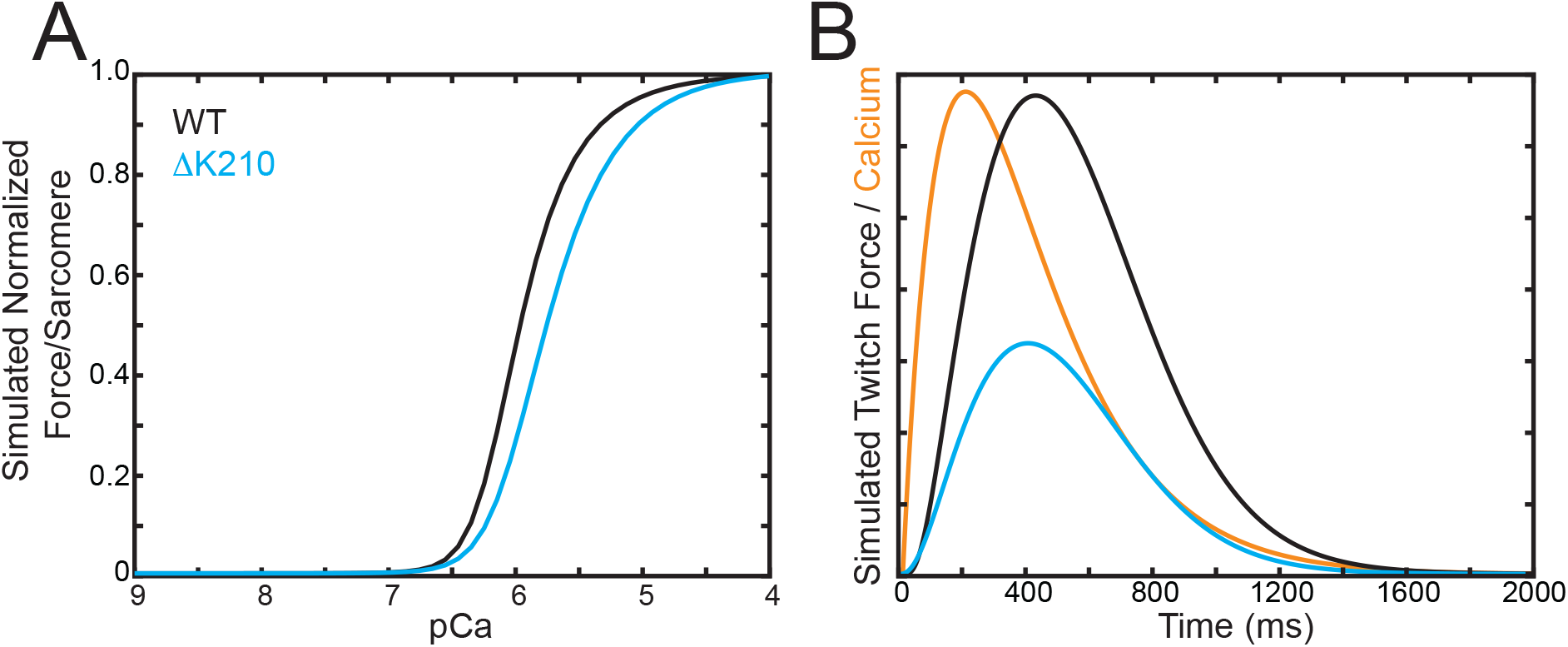
Computational modeling of the steady-state and transient force per sarcomere. (A) Using the model developed by Campbell et al. (52), the steady-state normalized force per sarcomere can be simulated for both ΔK210 and WT. The simulation shows a shift in the pCa50 towards supermaximal calcium activation in the mutant. (B) Simulated twitch forces per sarcomere in response to a calcium transient. The model predicts that ΔK210 should show a lower twitch force per sarcomere.

To predict the effects of the ΔK210 mutation on the force per sarcomere, we used the same computational model to simulate the force per sarcomere in response to a calcium transient. In the modeling, we assumed that the primary effects of the mutation are on thin filament positioning, and that to a first order approximation, the calcium transient is not significantly changed by the mutation. The consequences of these assumptions are explored in the Discussion. The simulation shows a smaller maximal twitch force per sarcomere in response to a calcium transient for ΔK210 compared to the WT (Figure 4B). Taken together, these simulations predict that ΔK210 should decrease the force per sarcomere in cardiomyocytes.

### Generation of ΔK210 induced stem cell-derived cardiomyocytes

To examine the effects of the ΔK210 mutation in human cells, we generated human induced pluripotent stem cell (hPSC)-derived cardiomyocytes (hPSC-CMs). The hPSC lines were derived from the BJ foreskin fibroblast line by the Washington University Genome Engineering core (see Materials and Methods). Whole exome sequencing of the stem cell line demonstrated that these cells do not have any genetic variants associated with familial cardiomyopathies (Figure 5 – figure supplement 1). Cell lines homozygous for the ΔK210 mutation were generated using the CRISPR/Cas9 system (17) (Figure 5 – figure supplement 2) (see Materials and Methods). ΔK210 mutant stem cells have normal karyotypes (Figure 5 – figure supplement 3) and are pluripotent, as assessed by immunofluorescence staining for pluripotency markers (Figure 5 – figure supplement 4). For all experiments with the mutant hPSC-CMs, two separate stem cell lines were used to ensure that the observations are not due to potential off-target cuts introduced during the CRISPR/Cas9 editing. We also used hPSC-CMs from at least two separate differentiations per cell line for both WT and ΔK210. hPSCs were differentiated to hPSC-CMs using established protocols (15, 53) involving temporal modulation of WNT signaling using small molecules (Figure 5 – figure supplement 5) and aged at least 30 days before use. Using this procedure, >90% cardiomyocytes were obtained, as determined by immunofluorescence staining for cardiac troponin-T (Figure 5 – figure supplement 5).

To examine the sarcomeric organization of hPSC-CMs, cells cultured on glass were fixed and stained for troponin-I and a-actinin to visualize the thin filament and z-discs, respectively (Figure 5). Unlike adult human tissue derived cardiomyocytes, which are rectangular with sarcomeres that align with the long axis of the cell, WT hPSC-CMs cultured on glass orient randomly and display robust sarcomeric staining that is typically not aligned along a single axis (Figure 5A). Compared to the WT hPSC-CMs, ΔK210 hPSC-CMs display more disorganized sarcomeres with patches of punctate staining (Figure 5B). Similar disorganization and punctate sarcomere structure has been seen with other hPSC-CM models of DCM cultured on stiff substrates (6, 18, 54), suggesting that it might be a common feature of some forms of DCM.

**Figure 5:**
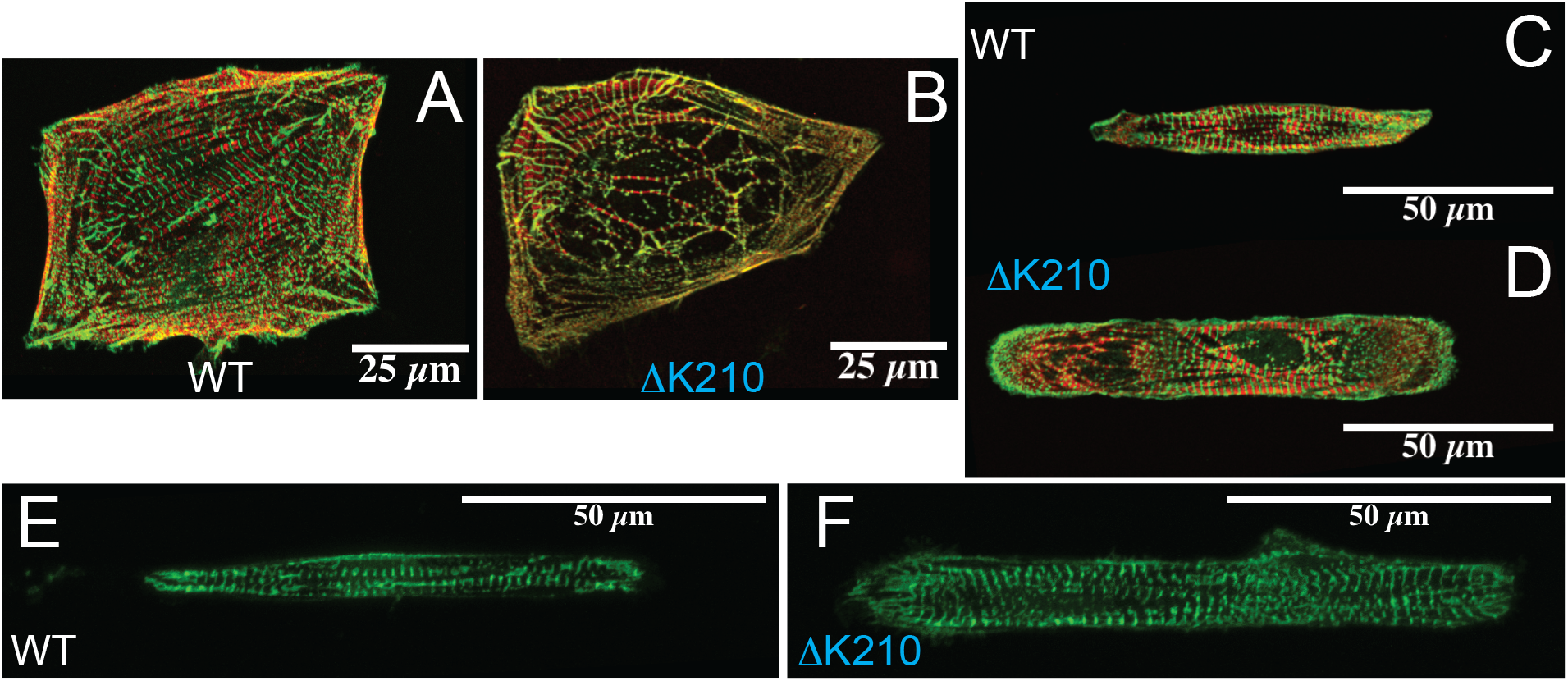
Immunofluorescence images of sarcomeres in hPSC-CMs. Troponin-I is red and a-actinin is green. All images are z-projections. (A) WT hPSC-CM on glass. (B) ΔK210 hPSC-CM on glass, showing sarcomeric disorganization. (C) WT cell on a rectangular pattern on glass. (D) ΔK210 cell on a rectangular pattern on glass. (E) WT cell on a rectangular pattern on a 10 kPa hydrogel. (F) ΔK210 cell on a rectangular pattern on a 10 kPa hydrogel. The sarcomeric organization is significantly improved on the hydrogel with physiologically relevant stiffness.

### ΔK210 hPSC-CMs show a pronounced increase in size and mechanosensitive alterations in sarcomeric structure

Cardiomyocytes are sensitive to their local physical environment, and it has been shown that providing physical cues that mimic the environment of the heart can improve cardiomyocyte structure and contractility (21, 55). During aging and DCM disease progression, the heart becomes stiffer, and the myocytes become disordered. We hypothesized that ΔK210 hPSC-CMs would show aberrant responses to changes in their physical environment. To test this hypothesis, we examined the size and structure of cells on glass, which has a high stiffness (~GPa) and on hydrogel substrates with a stiffness matched to healthy heart tissue (10 kPa).

To examine the role of geometric cues in cellular organization, we micropatterned extracellular matrix for cellular adhesion in rectangular patterns with a 7:1 aspect ratio (21). We examined live cells using bright field microscopy, and fixed cells stained for the z-disc marker a-actinin and troponin-I using confocal microscopy (Figure 5C-F). Interestingly, the ΔK210 hPSC-CMs are significantly larger than the WT in fixed cells on glass (Figures 5 and 6), fixed cells on 10 kPa hydrogels (Figures 5 and 6), and in live cells on 10 kPa hydrogels (Figures 5 and 7). We analyzed a large number of cells and generated cumulative distributions of cell sizes for single cells, since this methodology does not assume a form for the underlying distribution (Figures 6 and 7). The cumulative distribution shows the fraction of cells with an area less than or equal to the value of the x-axis. The increase in size is due to increases in both cell width and length (Table 1, Figure 6). These data demonstrate that ΔK210 hPSC-CMs are larger than the WT cells, and that this increase in size is not dependent on the mechanical environment.

**Table 1:** Analysis of Fixed Cell Sizes on Rectangular Patterns Patterned on glass

|  | **WT (n=120)** | **ΔK210 (n=66)** | **p** |
| --- | --- | --- | --- |
| **Area ($\mu\text{m}^2$)** | 1000 (-50/+50) | 1470 (-70/+70) | 0.00002 |
| **Length ($\mu\text{m}$)** | 91 (-3/+3) | 108 (-3/+3) | 0.00002 |
| **Width ($\mu\text{m}$)** | 13 (-0.4/+0.4) | 17 (-0.5/+0.5) | 0.00002 |
| **Length : width ratio** | 6.7 (-0.3/+0.3) | 6.4 (-0.2/+0.3) | 0.1 |
| **OOP** | 0.28 (-0.02/+0.02) | 0.24 (-0.02/+0.02) | 0.008 |

**Patterned on 10 kPa hydrogels**
|  | **WT (n=33)** | **ΔK210 (n=38)** | **p** |
| --- | --- | --- | --- |
| **Area ($\mu\text{m}^2$)** | 770 (-80/+90) | 990 (-60/+60) | 0.0002 |
| **Length ($\mu\text{m}$)** | 85 (-7/+7) | 96 (-4/+4) | 0.007 |
| **Width ($\mu\text{m}$)** | 11 (-1/+1) | 13 (-0.7/+0.8) | 0.06 |
| **Length : width ratio** | 7.6 (-0.9/+0.9) | 7.6 (-0.6/+0.6) | 0.9 |
| **OOP** | 0.28 (-0.04/+0.04) | 0.27 (-0.03/+0.03) | 0.5 |

**Figure 6:**
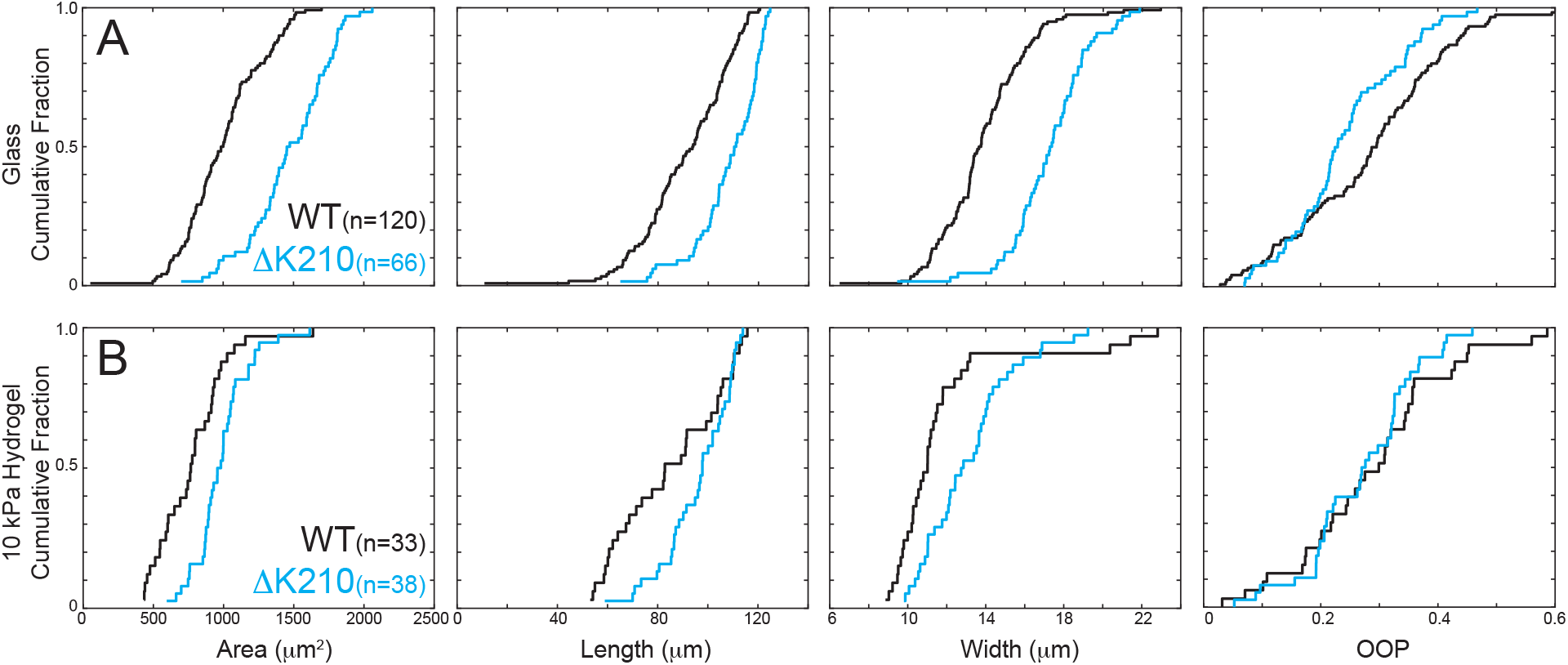
Quantification of immunofluorescence staining for sarcomeres in hPSC-CMs on rectangular patterns on (A) glass and (B) 10 kPa hydrogels. Data are plotted as cumulative distributions and parameter values can be found in Table 1. ΔK210 cells are significantly larger on both glass and 10 kPa hydrogels (p = 0.00002), due to increases in both length and width. The orientation order parameter (OOP) is a measurement of sarcomeric organization. For a hPSC-CM in which all of the sarcomeres are aligned along a single axis the OOP=1 and for a hPSC-CM with no preferred axis, the OOP = 0. On glass, ΔK210 cells have lower OOP values than the WT (p = 0.008). On physiological stiffness hydrogels, there is no significant difference between the OOP of the WT and mutant cells (p = 0.5).

**Figure 7:**
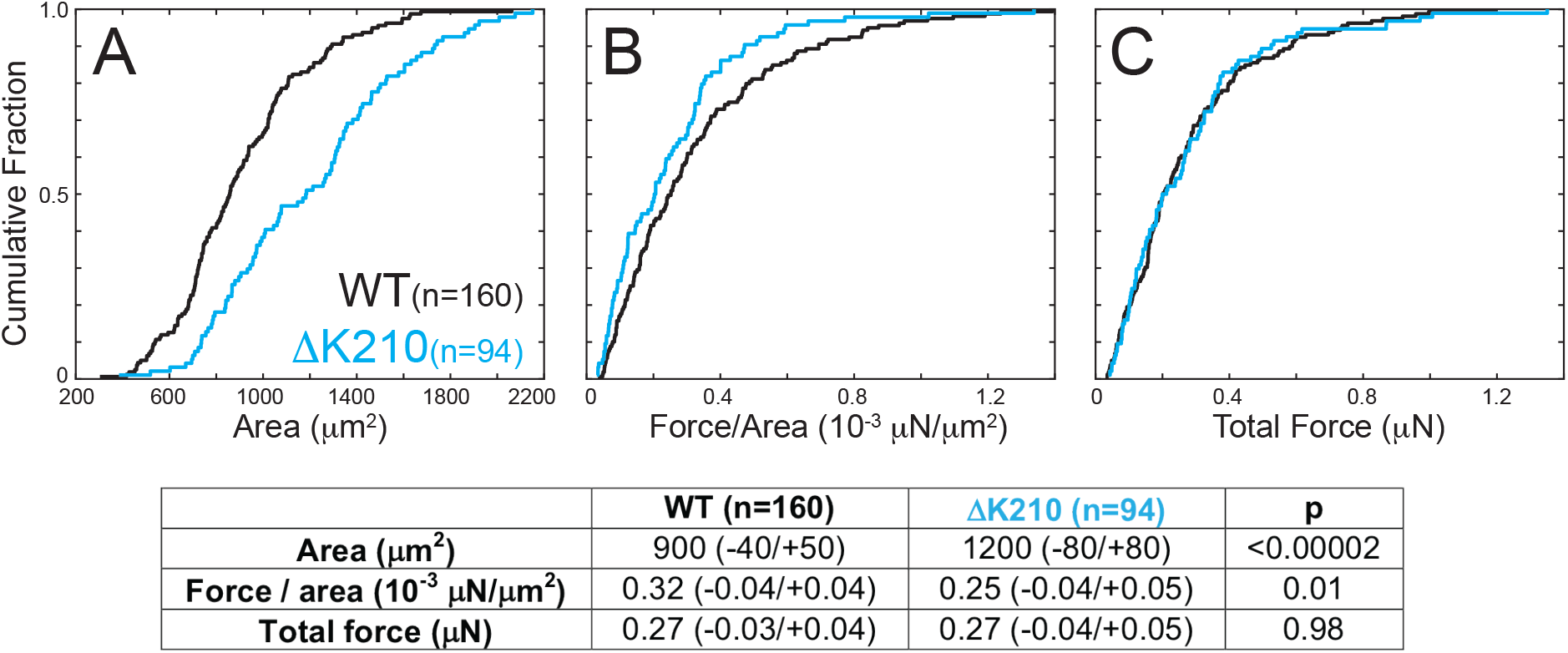
Traction force microscopy of single hPSC-CMs patterned on 10 kPa polyacrylamide gels. (A) Live ΔK210 hPSC-CMs on hydrogels are significantly larger than WT cells. (B) ΔK210 cells have a lower force per area (i.e., force per sarcomere), consistent with the molecular studies; however, since they are larger, (C) their total force production is not significantly different from the WT cells.

Next, we examined the influence of substrate stiffness on the sarcomeric organization of ΔK210 hPSC-CMs. We fixed and stained micropatterned hPSC-CMs for the sarcomeric marker a-actinin on both glass and 10 kPa substrates. Both WT and ΔK210 hPSC-CMs sarcomeres orient preferentially along the long axis of the cells patterned on glass and on 10 kPa hydrogels (Figure 5). To quantify the degree of sarcomeric organization, we used a program developed by Pasquilini et al. (55). The program analyzes fluorescence images with periodic structure, and it calculates the orientational order parameter, OOP (i.e., the fraction of z-discs that are oriented along a single axis). For cells with z-discs all oriented along a single axis, OOP=1, and for cells with z-discs with no preferred axis, OOP=0.

When patterned on glass (GPa stiffness), the ΔK210 mutant hPSC-CMs show a reduction in the fraction of sarcomeres aligned along the long axis compared to the WT (p = 0.008) (Figure 6A, Table 1). This is consistent with what we saw with unpatterned cells on glass (Figure 5A-B). Strikingly, when patterned on hydrogels that mimic the stiffness of healthy heart tissue, the ΔK210 cells show significantly improved sarcomeric organization (Figure 6B, Table 1). As quantified using the OOP, the organization of the ΔK210 sarcomeres are indistinguishable from the WT on 10 kPa substrates (p = 0.5). Taken together, these results demonstrate that the ΔK210 mutation affects how the hPSC-CMs respond to their mechanical environment, with ΔK210 cells showing normal sarcomeric organization on physiological stiffness substrates, but reduced sarcomeric organization on stiff substrates.

### ΔK210 cells show altered contractility and features of hypertrophy on substrates of physiological stiffness

Our computational modeling predicts that ΔK210 cells should show a lower force per sarcomere (Figure 4B); and therefore, we examined whether the ΔK210 mutation affects force production of single hPSC-CMs. To measure force production of single hPSC-CMs, we used traction force microscopy. Spontaneously beating cells were micropatterned onto 10 kPa hydrogels prepared with embedded fluorescent microbeads to monitor beating. Spinning disk confocal imaging was used to monitor the bead displacement as a function of time, and the total force of contraction for each cell was calculated using a MATLAB routine developed by Ribeiro et al. (56) (Figure 7B-C). For each cell, the force per sarcomere was approximated by calculating the force per area.

The average total force per cell measured by traction force microscopy was the same for both WT (0.27 (−0.03/+0.04) μN) and ΔK210 hPSC-CMs (0.27 (−0.04/+0.05) μN; p = 0.98) (Figure 7C). For each cell analyzed by traction force microscopy, we measured its area from bright-field illumination images (Figure 7A). Bright-field images of live cells lack the resolution of the immunofluorescence images, making it impossible to accurately measure the cross-sectional area of the live cells; however, ΔK210 hPSC-CMs are significantly larger than the WT cells (Figure 7A). Moreover, fixed ΔK210 hPSC-CMs on 10 kPa hydrogels have a sarcomeric organization that is indistinguishable from the WT (Figure 6B). We calculated the force per area for each cell (Figure 7B). The ΔK210 cells have a lower force per area (0.25 (−0.04/+0.05) μN/μm^2^) compared to the WT (0.32 (−0.04/+0.04) μN/μm^2^; p = 0.01). The lower force per area for ΔK210 is consistent with our molecular studies and the computational prediction of a lower force per sarcomere (Figure 4B). These data link our molecular and cellular phenotypes. Moreover, they demonstrate that although the force per sarcomere is reduced in the mutant, the ΔK210 hPSC-CMs can compensate through an increase in size that results in a total force per cell that is indistinguishable from the WT.

## Discussion

Here, we determined the mechanism of a mutation in troponin-T that causes DCM in humans, ΔK210. We found that this mutation affects not only molecular and cellular contractility, but also the ability of the cardiomyocytes to respond to changes in the mechanical environment like stiffening of heart tissue associated with aging and disease in patients. These results implicate defective mechanosensing as an important factor in the pathogenesis of DCM caused by mutations in sarcomeric proteins, and they highlight the importance of multiscale studies in understanding heart disease.

### Proposed molecular mechanism for ΔK210 and its relationship to previous studies

The disease presentation of familial cardiomyopathies and their biophysical manifestation (i.e., contractile and force-dependent properties) can depend on the myosin isoform (7, 8, 57–60). Our *in vitro* motility results (Figure 1B) using porcine cardiac actin and ventricular β-cardiac muscle myosin (MYH7), which closely mimics the biophysical and biochemical properties of human ventricular β-cardiac muscle myosin (38, 39), demonstrate a shift towards supermaximal calcium activation, consistent with previous experiments studying the ΔK210 using different myosin isoforms (31, 44).

We determined that this shift is due to alterations in the equilibrium constants that govern the positioning of tropomyosin along the thin filament, leading to a reduction in the population of force-generating cross bridges at submaximal calcium concentrations (Figures 3 and 4A). The structural basis of these mutation-induced shifts is not well understood; however, it is possible that the mutation affects the interaction between the troponin complex and tropomyosin (61) and/or the allosteric coupling between subunits of the troponin complex (2, 5, 62, 63). These non-exclusive mechanisms could contribute to the observed decrease in force-producing states seen in ΔK210 thin filaments, and future studies should be able to shed light on the structural basis of these changes.

Our data were analyzed using the formalism developed by McKillop and Geeves (47). Other models of thin filament regulation exist (64–68), but the McKillop and Geeves model is one of the most frequently used. Due to the nature of our experimentation, it is not possible to directly compare our results with those from other models of thin filament regulation. However, the computational simulations based on the McKillop and Geeves model are able to recapitulate the shift in calcium sensitivity seen in the motility assay (Figures 1B and 4A).

The computational modeling based on our biochemical studies predicts a larger reduction in the force per sarcomere than we observed in cells (Figure 7). This is likely due to several necessary simplifying assumptions made in the modeling: 1) We assumed that the mutation does not change the magnitude or time course of the calcium transient, which is not necessarily true given that the knock-in ΔK210 mouse shows altered calcium transients (31). 2) We assumed that the troponin in hPSC-CMs has the same biochemical properties as the adult cardiac troponin complex that we examined in our biochemical experiments. This is not necessarily the case, since four troponin-T isoforms can be detected in the human heart at different developmental stages (69, 70), and hPSC-CMs primarily express a slow skeletal muscle isoform of troponin-I (71), rather than the cardiac isoform used in the biochemical studies. While all isoforms of troponin expressed in hPSC-CMs would have the ΔK210 mutation (Figure 5 – figure supplement 2), different isoforms have different calcium sensitivities (69). That said, even with these simplifying assumptions, the simulations were able to predict the reduction in the force per sarcomere seen in the ΔK210 cardiomyocytes.

### Linking the molecular and cellular contractile phenotypes for ΔK210

Patients with the ΔK210 mutation show early onset of the disease phenotype and a high incidence of sudden death (26). Aspects of the disease have been replicated in knock-in mouse models (31), which have significantly advanced our understanding of the disease phenotype. However, the ΔK210 mouse model shows significant differences with respect to the human phenotype, due to inherent physiological differences between mouse and human hearts, including differences in gene expression, calcium-handling machinery, and ion channel composition (34).

We used human hPSC-CMs to study how the initial insult of a molecular-based change leads to the early disease pathogenesis in human cells. For these studies, hPSC-CMs are excellent models, since they can capture cellular changes that occur before major compensatory mechanisms (e.g., fibrosis, activation of neurohormonal pathways, changes in gene expression) mask the initial molecular phenotype (18–20, 54, 72). Contractile changes associated with the disease have been observed in cardiomyopathy patients before adverse remodeling of the heart (73–75). That being said, hPSC-CMs are developmentally immature, meaning that they do not necessarily recapitulate all the aspects of the phenotype in adults or the later stages of the disease progression. All of our molecular and cellular studies modeled the homozygous ΔK210 mutation, so care should be used when extrapolating the results here to the phenotype seen in patients.

ΔK210 hPSC-CMs show a reduced force per area compared to WT cells (Figure 7B). We did not see any differences in the sarcomeric organization of mutant cells on patterned hydrogels (Figure 6B), suggesting that the observed reduction in the force per area is due to a reduction in the force per sarcomere. This result is consistent with the results of our computational modeling of our biochemical measurements (Figure 4B), and it links the molecular phenotype with cellular contractility.

Interestingly, while the force per sarcomere is reduced in the mutant, the total force per cell is not different from the WT (Figure 7C). This can be explained by the fact that the mutant cells are larger (Figures 6, 7), a result that would not have been predicted from the molecular studies alone. This increase in size occurs in single cells, independent of endocrine or cell-cell signaling. One possible explanation for this increase in size is that the cells are undergoing compensatory hypertrophy, which allows the cells to counteract the reduced force per sarcomere. While we have not investigated whether the increase in size seen in our cells is due to activation of hypertrophic signaling, future studies should elucidate the mechanism of the increase in cell size seen here.

### Defects in ΔK210 mechanosensing contribute to the disease phenotype

Fundamentally, troponin serves as a gate that regulates myosin-based tension. As such, one would expect that the primary effects of the mutation would be to reduce force production; however, the ΔK210 cells show several changes beyond altered contractility. While healthy cardiomyocytes adapt their contractility and structure to changes in their mechanical environment, such as age or disease-related stiffening of the heart tissue (21), ΔK210 cells do not adapt in the same way as the WT cells. Similar to other hPSC-CM models of DCM grown on stiff substrates, ΔK210 hPSC-CMs cultured on glass show impaired sarcomeric organization (18, 54, 76); however, ΔK210 hPSC-CMs can organize normally when given mechanobiological cues that mimic the healthy heart (Figures 5 and 6). Therefore, the ΔK210 mutation affects not only contractility, but also the ability of the hPSC-CMs to sense and respond to their environment.

The connection between the reduction in myosin-generated tension in the mutant cells and an impaired ability to respond to mechanical forces warrants further investigation; however, a possible link between cellular tension and sarcomeric organization in DCM was revealed by live-cell imaging of sarcomerogenesis in hPSC-CMs (54). It was shown that pharmacological inhibition of myosin contractility or genetic disruption of titin leads to an impaired ability of cells to generate well-organized sarcomeres. Chopra et al. proposed that the transduction of myosin-driven tension is necessary for the proper assembly of sarcomeres. We propose that a similar mechanism could be relevant in ΔK210 cells, which show reductions in myosin-based contractility and sarcomeric disorganization when patterned on glass. Consistent with this idea, treatment of hPSC-CMs containing a different DCM mutation with Omecamtiv mecarbil, a myosin-based thin filament activator (41, 77, 78), leads to an improvement in sarcomeric organization (76). We speculate that some of the salutary effects of the drug are due to the restoration of tension necessary for proper mechanosensing.

The result that ΔK210 hPSC-CMs show altered responses to changes in their mechanical environment has important implications for the mechanism of the disease pathogenesis. Previous studies of cardiomyocytes have shown that the organization of sarcomeres varies depending on substrate stiffness (21, 76), with peak contractility occurring on substrates of physiological stiffness. ΔK210 hPSC-CMs can organize properly and they have normal force production on substrates that match the stiffness of the healthy heart (Figures 6 and 7); however, heart tissue becomes stiffer with the disease progression and aging. We propose that as the heart tissue becomes stiffer, ΔK210 cells would become more disordered, effectively causing a progressive loss of function, and potentially contributing to myocyte disarray. Thus, the inability of the mutant cells to adapt to changes in their mechanical environment may contribute to the disease pathogenesis and progression.

### The potential importance of mechanosensing in DCM pathogenesis

Our results suggest that reduced force at the molecular scale can lead to impaired mechanosensing in cardiomyocytes. While the disease presentation in DCM depends on the exact mutation (79), we propose that disruption of mechanosensing could be a common mechanism in the disease pathogenesis of other sarcomeric and non-sarcomeric DCM mutations. It has been proposed that DCM can be caused by molecular hypocontractility (4, 80, 81); however, there are also many DCM-causing mutations in non-sarcomeric genes with no known roles in contractility; suggesting that other mechanisms might be involved (1, 3, 82). Most, but not all, of these genes are located along proposed mechanosensing pathways (83, 84) used to sense and transduce mechanical forces. For example, some of these genes link the cytoskeleton to the extracellular matrix (e.g., dystrophin (DMD), vinculin (VCL)), others link the cytoskeleton to the nucleus (e.g., nesprin-1 (SYNE1), lamin-A (LMNA), centromere protein F (CENP-F)), and others are mechanosensitive transcription factors (e.g., TAZ). Consistent with this idea, altered mechanobiology has been identified in DCM cells with mutations in the non-sarcomeric protein lamin (82). Moreover, patients with mutations in these genes, such as those with Duchenne muscular dystrophy and Emery-Dreifuss muscular dystrophy, often develop DCM. It is intriguing to speculate that impaired mechanosensing by cardiomyocytes in the heart could link some of these seemingly different diseases.

## Conclusions

Using a combination of biochemical, computational, and cell biological techniques, we reveal several features of a DCM mutation that would have been missed by studying the molecular or cellular phenotypes alone. We demonstrate that the ΔK210 mutation in human cardiac troponin-T causes a change in the equilibrium positioning of tropomyosin, leading to alterations in the calcium sensitivity of thin filament activation. Computational modeling predicted that these changes should lead to a reduction in the force generated per sarcomere in mutant cells, and we demonstrate that this is indeed the case in hPSC-CMs. Moreover, our results demonstrate that the ΔK210 mutation affects the structural organization of hPSC-CMs and alters how they respond to their mechanical environment. Finally, we demonstrate that the ΔK210 hPSC-CMs can increase in size to normalize their force production. Taken together, these results demonstrate that disease-causing mutations of sarcomeric proteins affect not only contraction, but also how cardiomyocytes sense and respond to changes in their mechanical environment associated with aging and disease. These data also suggest that disrupting mechanosensing pathways can contribute to the disease phenotype in DCM.

## Acknowledgements

We would like to thank Stuart Campbell and Francesco Pasqualini for sharing their code for the simulations and sarcomeric structure analysis, respectively. We acknowledge Mike Ostap and Ken Margulies for extremely helpful discussions during the early planning phases of this project. We would also like to acknowledge Drew Braet for technical assistance and Samantha Barrick for assistance with biochemical experiments. Funding for this project was provided by a pilot grant from the Children’s Discovery Institute of Washington University and St. Louis Children’s Hospital, the Washington University Center for Cellular Imaging (WUCCI) (CDI-CORE-2015-505), the National Institutes of Health (R00HL123623, R01HL141086 to M.J.G., T32EB018266 to S.R.C.), and the March of Dimes Foundation (FY18-BOC-430198 to M.J.G.).

## Author contributions

S.R.C. purified proteins, performed and analyzed the stopped flow and fluorescence experiments, and drafted the manuscript. P.E.C. performed and analyzed the fluorescence and traction force microscopy experiments with the stem cell derived cardiomyocytes. L.G. purified proteins, performed *in vitro* motility assays, implemented the cell-based assays, and performed and analyzed experiments with stem cell derived cardiomyocytes. M.E. performed and analyzed fluorescence microscopy experiments. W.T.S. designed tools for microcontact printing. M.J.G. oversaw the project, generated mutant proteins, implemented biochemical assays, analyzed data, and drafted the manuscript. All authors contributed to the editing of the final manuscript.

## Materials and Methods

### Tissue purification of proteins for *in vitro* biochemical experiments

Porcine cardiac myosin and actin were simultaneously purified from cryoground porcine ventricles (Pelfreez) as previously described (38). Myosin subfragment-1 (S1) was prepared by chymotrypsic digestion of myosin using the method of Margossian and Lowey (85) with the modifications of Eads et al. (86). Purity of S1 was assessed by SDS-PAGE, and the protein concentration was determined using the Bradford reagent (Thermo Scientific). Porcine cardiac actin was purified from acetone powder (87) and labeled with N-(1-pyrenyl)iodoacetamide (pyrene) as described previously (88, 89). Actin concentrations were determined by absorbance at 290 nm (and 344 nm for pyrene-labeled actin).

### Preparation of recombinant human cardiac tropomyosin and troponin

pET3d vectors containing human cardiac tropomyosin, troponin-I, troponin-T, and troponin-C were a generous gift from L. Tobacmann (University of Illinois at Chicago). The ΔK210 mutation was introduced into troponin-T using a QuickChange Site-Directed Mutagenesis Kit (Agilent) and verified by sequencing. Human cardiac troponin complex subunits were expressed in *E. coli* and purified from BL21-CodonPlus (Agilent) using established protocols (90), with the modification that the final protein was purified over a MonoQ column (GE Healthcare) instead of a ResourceQ column. Troponin complex concentration was measured using the Bradford reagent. Recombinant human tropomyosin was purified from BL21-CodonPlus (Agilent) using established protocols (91) with the modifications of McIntosh et al. (92). Tropomyosin concentration was determined using absorbance at 280 nm. Before use in biochemical assays, tropomyosin was reduced in 50 mM DTT at 56 °C for 5 minutes, and then aggregates were removed by ultracentrifugation (92).

### *In vitro* motility assay

*In vitro* motility assays were conducted as previously described (36). Phalloidin-stabilized F-actin and rhodamine-phalloidin-stabilized F-actin were each prepared in KMg25 buffer (25 mM KCl, 2 mM EGTA, 60 mM MOPS, 1 mM DTT, and 4 mM free MgCl_2_). To remove myosin dead-heads, full-length cardiac myosin in high salt buffer (KMg25 with 300 mM KCl) with 2.17 μM phalloidin-stabilized F-actin and 1 mM ATP was sedimented at 355040 x g for 30 minutes at 4°C. A Bradford assay was used to determine the concentration of functional myosin heads following sedimentation. Flow cells were assembled with two glass coverslips (one coated with nitrocellulose) using double stick tape and vacuum grease. One volume of myosin (200 nM) was added to the flow cell followed by 1 volume of 1 mg/mL BSA, 1 volume unlabeled actin, 2 volumes of KMg25 + 1 mM ATP, 4 volumes of KMg25, and then 1 volume of reconstituted thin filaments (40 nM rhodamine-phalloidin-labeled actin with 2 μM troponin and tropomyosin). One volume equals 50 μL, which is slightly larger than the flow cell volume. Activation of motility was initiated by adding 2 volumes of activation buffer (KMg25 with 4 mM ATP, 1 mg/mL glucose, 192 U/mL glucose oxidase, 48 μg/mL catalase, 2 μM troponin and tropomyosin, 0.5% methyl cellulose). The speed of thin filament translocation was recorded at room temperature (~20 °C) over 20 frames. 25 consistently moving filaments from each video were manually tracked and analyzed using the MTrackJ plugin on Fiji ImageJ (93). Data were collected from 3 separate experiments.

### Stopped-flow transient kinetics to determine *K*_B_ and the rate of ADP release

To determine *K*_B_, the rate of myosin binding to regulated thin filaments was measured at 20 °C using stopped-flow techniques (47) in an SX-20 apparatus (Applied Photophysics). Pyrene-actin was excited at 365 nm, and fluorescence emission was detected using a 395 nm long-pass filter. 5 μM phalloidin-stabilized pyrene actin, 2 μM tropomyosin, 2 μM troponin, and 0.04 U/mL apyrase were rapidly mixed with 0.5 μM S1 myosin and 0.04 U/mL apyrase. The high calcium (pCa 4) buffer contained 200 mM KCl, 5mM MgCl_2_, 60 mM MOPS, 2 mM EGTA, 1mM DTT, and 2.15 mM CaCl_2_. The low calcium (pCa 9) buffer contained 200 mM KCl, 5mM MgCl_2_, 60 mM MOPS, 2 mM EGTA, 1 mM DTT and 5.2 μM CaCl_2_. Each experiment was the average of at least 3 separate mixes, and the data were fit by a single exponential curve. *K*_B_ was calculated by using Equation 1. The average *K*_B_ was calculated from three different experiments and the p-value was calculated using a 2-tailed Student’s t-test.

The rate of ADP release from myosin bound to regulated thin filaments at 20 °C was determined using stopped-flow techniques (94). 1 μM phalloidin-stabilized pyrene-actin, 1 μM S1 myosin, 100 μM ADP, 100 μM MgCl_2_, 2 μM tropomyosin, and 2 μM troponin were rapidly mixed with 5 mM ATP and 5 mM MgCl_2_. The experiment was performed in high calcium (pCa 4) buffer containing 25 mM KCl, 5mM MgCl_2_, 60 mM MOPS, 2 mM EGTA, 1 mM DTT, and 2.15 mM CaCl_2_. The rate of ADP release was calculated by fitting a single exponential function to the data, as we have previously done (89) (Figure 1 – figure supplement 1). The average of three experiments was calculated, and statistical testing was done using a two-tailed Student’s t-test.

### Fluorescence titrations to determine the values of *K*_W_, *K*_T_, and *n*

Fluorescence titrations were carried out at 20 °C as described in (47) to determine the values of *K*_W_, *K*_T_, and *n*. S1 myosin (up to a final concentration of 10 μM) was added at 1-minute intervals to a stirred cuvette containing 0.5 μM pyrene-actin and 0.27 μM troponin and tropomyosin. Titrations were performed at low (2 mM EGTA), intermediate (pCa 6.25) and high (pCa 3) calcium concentrations. The buffers contained 200 mM KCl, 5 mM free MgCl_2_, 60 mM MOPS, 2 mM EGTA, 1 mM DTT, 2 mM ADP, and the desired free concentration of calcium from CaCl_2_. The buffer also contained 50 μM P1,P5-di(adenosine-5’)pentaphosphate (Ap5A), 2 mM glucose, and 1 μM hexokinase to eliminate any contaminating ATP. Five repeats were performed for each condition. Titration curves were globally fit using an annealing algorithm in MATLAB (MathWorks) (50). 95% confidence intervals were determined by fitting 1000 bootstrap simulations. A test statistic was defined by calculating the difference in fit parameters between bootstrap simulations of the WT and mutant data, and the p-value was determined from the fraction of test statistic simulations that did not include the null hypothesis (i.e., that the parameters for the WT are the same as the mutant). The fraction of thin filament in each equilibrium state (Figure 3B) was calculated using the partition function.

### Simulations

Simulations of force production by sarcomeres were conducted using the model developed by Campbell et al. (52). This model enables the calculation of the normalized force-pCa curve and the twitch force in response to a calcium transient for a given set of equilibrium constants. Code for the simulation was kindly provided by Stuart Campbell. For simulation of the WT protein, the default parameters were used, except the value of the forward rate constant out of the blocked state was adjusted to equal 0.3 to capture the measured value of *K*_B_. This same *K*_B_ value was used for the mutant simulation, since Kb was unchanged in the mutant. To simulate the mutant, the forward rate constant from the closed to open state was set to 8.35 so that the equilibrium constant would be proportional to the measured *K*_T_.

### Stem cell line derivation

The parent stem cell line, BJFF.6, was generated from the human BJ fibroblast line (CRL-2522, ATCC) by the Genome Engineering and iPSC Center (GEiC) at Washington University in St. Louis. hPSC reprogramming was done using the CytoTune-iPS 2.0 Sendai reprogramming kit (A16517, ThermoFisher) following the manufacturer’s recommended protocol.

### Whole exome sequencing

Whole exome sequencing of the BJFF stem cell line was performed by the McDonnell Genome Institute (Washington University in St. Louis). DNA was isolated from a frozen cell pellet using the QiaAMP DNA Mini Kit (Qiagen) according to the manufacturer’s recommendations. DNA was quantified fluorometrically using the Qubit and dsDNA HS Assay kit (Invitrogen). The DNA yield was 22 μg / 5 million cells.

An automated dual-indexed library was constructed with 250 ng of genomic DNA utilizing the KAPA HTP Library Kit (KAPA Biosystems) on the SciClone NGS instrument (Perkin Elmer) targeting 250 bp inserts. 1 μg of library was hybridized with the xGen Exome Research Panel v1.0 reagent (IDT Technologies) that spans 39 Mb target regions (19,396 genes) of the human genome. The concentration of the captured library pool was accurately determined through qPCR (Kapa Biosystems) to produce cluster counts appropriate for the S2 flow cell on the NovaSeq platform (Illumina). A 2 x 151 base pair end sequence data generated approximately 15 Gb per sample which led to greater than 90% of the targets covered at a minimum of 20x depth of coverage. The mean depth of coverage was approximately 200x. Variants were identified using the Variant Effect Predictor. Variants in genes that have been associated with cardiomyopathies (ABCC9, ACTC1, ACTN2, ALMS1, ALPK3, ANKRD1, BAG3, BRAF, CAV3, CHRM2, CRYAB, CSRP3, DES, DMD, DOLK, DSC2, DSG2, DSP, DTNA, EMD, FHL1, FKRP, FKTN, FLNC, GATAD1, GLA, HCN4, HRAS, ILK, JPH2, JUP, KRAS, LAMA4, LAMP2, LDB3, LMNA, MAP2K1, MAP2K2, MIB1, MTND1, MTND5, MTND6, MTTD, MTTG, MTTH, MTTI, MTTK, MTTL1, MTTL2, MTTM, MTTQ, MTTS1, MTTS2, MURC, MYBPC3, MYH6, MYH7, MYL2, MYL3, MYLK2, MYOZ2, MYPN, NEBL, NEXN, NKX2-5, NRAS, PDLIM3, PKP2, PLN, PRDM16, PRKAG2, PTPN11, RAF1, RBM20, RIT1, RYR2, SCN5A, SGCD, SOS1, TAZ, TCAP, TGFB3, TMEM43, TMPO, TNNC1, TNNI3, TNNT2, TPM1, TTN, TTR, TXNRD2, VCL) were cross referenced with the ClinVar database. None of the variants identified in the BJFF cell line have been characterized as pathogenic or likely pathogenic in patients (Figure 5 – figure supplement 1).

### Derivation of the ΔK210 stem cell lines

Two independent stem cell lines homozygous for the ΔK210 deletion, AN184.2-hTNNT2, were generated from the BJFF.6 human hPSC line by Genome Engineering and iPSC Center (GEiC) at Washington University in St. Louis using the CRISPR/Cas9 system (17, 95). The oligo used to generate the gRNA was ACACCGAGAAGATTCTGGCTGAGAGGG (Figure 5 – figure supplement 2). This gRNA was selected to minimize potential off-target effects based on an analysis of similar sequences. The mutation was introduced using homology-directed repair to a singlestranded oligodeoxynucleotide template (ssODN) with the deletion. Approximately 1 to 1.5 × 10^6^ hPSCs were washed in DPBS and resuspended in P3 primary buffer (Lonza) with 1 μg gRNA, 1.5 μg Cas9 vectors, and 1 μL ssODN (4μg, IDT) and then electroporated with a 4D-Nucleofector (Lonza) using the CA-137 program. Following nucleofection, cells were single-cell sorted (96) and screened with targeted deep sequencing analysis (97) using primer sets specific to target regions. The ΔK210 deletion was found in ~10% of the clones. Two clones with the ΔK210 mutation were then used for all cellular experiments.

After gene editing, karyotype (G-banding) analysis was performed by Cell Line Genetics. Cytogenetic analysis was performed on 20 G-banded metaphase cells (Cell Line Genetics) and these cells had normal karyotypes (Figure 5 – figure supplement 3). Cells were tested for myoplasma infections by the Genome Engineering core.

Pluripotency of gene-edited hPSCs was characterized by staining for SSEA4, OCT4, SOX2, and TRA-1-61 using the Pluripotent Stem Cell 4-Marker Immunocytochemistry kit (A25526, ThermoFisher) following the manufacturer’s recommended protocol. Immunofluorescent images of the stained cells were captured using a Nikon fluorescence microscope and CCD camera. Cells showed robust staining for pluripotency markers (Figure 5 – figure supplement 4).

### Stem cell culture and differentiation to hPSC-CMs

All culture media contained 10 U/mL penicillin and 10 μg/mL streptomycin. hPSCs were maintained at 37 °C and 5% CO2 in feeder-free culture on Matrigel (Corning) coated plates in Essential 8 Flex or StemFlex media (ThermoFisher). Cells were passaged using 0.02% EDTA (Sigma). After passaging, the E8 media was supplemented with 5 μM of the ROCK inhibitor Y-27632 (Selleck Chem).

Differentiation to cardiomyocytes was accomplished using the method of Lian et al. (53) (Figure 5 – figure supplement 5A). Briefly, hPSC colonies were dissociated to single cells using Accutase (Gibco) and seeded into monolayers on 12-well Matrigel coated plates at a density of 0.5-1 million cells per well. Cells were maintained in StemFlex media supplemented with 5 μM Y-27632 for the first 24 hours. Two days after seeding (day 0), the media was changed to RPMI/B27-I (RPMI-1640 media (Sigma) containing B-27 supplement without insulin (ThermoFisher). 6-8 μM of the WNT agonist CHIR-99021 (Selleck Chem) was added to initiate differentiation for 24 hours along with 1 μg/mL insulin to improve cell survival. On day 1, the WNT agonist was removed and cells were cultured in RPMI/B27-I. On day 3, the media was collected and centrifuged to remove debris. The conditioned media was combined with fresh RPMI/B27-I media in a 1:1 ratio and was supplemented with 5 μM of the WNT inhibitor IWP-2. On day 5, the IWP2 was removed and the cells were cultured in RPMI/B27-I. On day 7, the media was changed to RPMI/B27+I (RPMI-1640 media containing B-27 supplement with insulin) and changed every 3 days. Spontaneous beating began on day 9-12.

To enrich the population of cardiomyocytes, metabolic selection was used (98). On days 12-17, the cells were cultured in glucose-free media (glucose-free RPMI-1640 supplemented with B-27 supplement with insulin). On day 19, the cells were dissociated using 0.25% trypsin and then passaged to gelatin coated culture dishes in RPMI-20 (RPMI-1640 + 20% FBS) with the addition of 10 μM Y-27632. Starting on day 21, cells were cultured in RPMI/B27+I and the media was changed every 3 days. Metabolic selection gave populations of >90% cardiomyocytes (Figure 5 – figure supplement 5B). Cells were aged to at least day 30 after differentiation before being used for assays.

### Fabrication of stamps for microcontact patterning

Stamps for microcontact printing were generated using standard microfabrication techniques, based on the procedure in Ribeiro et al. (21). A photomask containing an array of 17 x 118 μm (total area of 2000 μm^2^) was generated (CAD/Art Services). SU-8-3010 (Microchem) was spin coated on to a 3” silicon wafer (University Wafer) using a Brewer Science CEE 200X instrument (Washington University Institute of Material Sciences and Engineering) to generate a 15 μm thick layer. The wafer was then soft baked at 65 °C for 90 seconds followed by 6 minutes at 95 °C. The mask exposure was done using a Karl Suss MJB 3UV 400 Mask Aligner for a total exposure energy of 200 mJ/cm^2^. Post exposure, the wafer was baked at 65 °C for 1 minute followed by 3 minutes at 95 °C. The un-polymerized SU-8-3010 was then removed from the master by washing in SU-8 developer (Y020100, Microchem).

Polydimethylsiloxane (PDMS) stamps for microcontact printing were prepared from the SU-8 master. The SU-8 master was silanized using vapor deposition of Trichloro(1H,1H,2H,2H-perfluoro-octyl) silane (Sigma) under vacuum. PDMS (Sylgard 184, Dow Corning) was prepared using a 10:1 w/w mixture of elastomer base:curing agent. PDMS was degassed under vacuum, poured on the master, and cured overnight under vacuum. The following day, the PDMS stamps were hard cured at 80 °C for 1 hour, peeled from the master, and then cut into 18 x 18 mm squares using a razor blade.

### Microcontact patterning of cardiomyocytes on glass

For patterning of cardiomyocytes on glass, glass coverslips were first cleaned by sonication in isopropanol. The glass was washed twice with water and dried using compressed nitrogen gas. The glass was then plasma cleaned under vacuum for 1 minute. The glass surface was passivated for 30 minutes in 0.1 mg/mL PLL-g-PEG (SuSoS AG), then washed with water, and dried with nitrogen gas. PDMS stamps were sterilized under UV light before use for at least 10 minutes. Geltrex (ThermoFisher) was diluted 1:10 in DMEM-F12 and then 40 μL was added to the surface of the PDMS stamp for 20 minutes. The Geltrex-coated PDMS stamp was dried with nitrogen gas and then placed Geltrex side down on the PLL-g-PEG treated coverslip. A 50 or 100 g weight was placed on top of the PDMS stamp for 2-3 minutes to transfer the Geltrex pattern onto the coverslip. The stamped glass coverslips were then transferred to the bottom of a 6-well tissue culture plate.

Cardiomyocytes were dissociated and singularized using 0.25% trypsin (Gibco). Cells were resuspended in RPMI-20 with 10 μM Y-27632 at a density of 500 cells/μL and then incubated on the patterned glass for 45 minutes. 18 mm coverslips required 300 μL volume per coverslip. The patterned cardiomyocytes were cultured in RPMI-20 with Rock inhibitor for 48 hours and were subsequently cultured in RPMI/B27+I with media changes every 3 days.

### Immunostaining and fluorescence microscopy

Cardiomyocytes were fixed for 15 minutes in 4% formaldehyde (Electron Microscopy Sciences) in phosphate-buffered saline (PBS) and permeabilized with 0.1% Triton X-100 (Sigma) for 20 minutes. The cells were blocked for 1 hour using a blocking solution containing 3% bovine serum albumin (Gold Bio), 5% donkey serum (Sigma), 0.1% Triton X-100, and 0.02% sodium azide in PBS. Primary antibodies (see table below) were added for 1 hour at room temperature or overnight at 4 °C. Cells were then washed thoroughly in PBS before incubating for 1 hour in secondary antibody. Cells were again thoroughly washed in PBS. For experiments with 4’,6-diamidino-2-phenylindole (DAPI), the stain was used at a 1:50,000 dilution. For experiments using tetramethylrhodamine B isothiocyanate labeled phalloidin, the phalloidin was added using a 1:1000 dilution. Cells were visualized using a Nikon A1Rsi confocal microscope or a Nikon spinning disk confocal microscope equipped with a Yokagawa CSU-X1 variable speed Nipkow spinning disk scan head and an Andor Zyla 4.2 Megapixel sCMOS camera (Washington University Center for Cellular Imaging). Z-stacks of cells with 60x magnification were recorded in sequential scanning mode.

### Antibodies used

| Antibody | Product Number | Working Dilution | Host species | Vendor |
| --- | --- | --- | --- | --- |
| Alpha-actinin | A7811/EA-53 | 1:500 | mouse | Sigma |
| Troponin-I | SC-15368 | 1:300 | rabbit | Santa Cruz |
| Cy3 anti-rabbit | 711165152 | 1:300 | donkey | Jackson Immuno. |
| Cy2 anti-mouse | 715225150 | 1:300 | donkey | Jackson Immuno. |
| OCT4 | A25532 | 1:400 | Rabbit | Invitrogen |
| SSEA4 | A25530 | 1:100 | Mouse | Invitrogen |
| SOX2 | A24759 | 1:100 | Rat | Invitrogen |
| TRA-1-60 | A24868 | 1:100 | Mouse | Invitrogen |

### Analysis of immunofluorescence images

Analysis of the organization of the sarcomeres was analyzed using software developed by the Parker lab (55). Standard deviation projections were generated from Z-stacks of hPSC-CMs stained for alpha-actinin. Data were pre-processed in FIJI/ImageJ (93) using an ImageJ plugin (55). Data were then analyzed using the Parker software to extract the orientational order parameter (OOP, a measure of sarcomeric alignment along a single axis) (55). The cellular area, length, and width were calculated from thresholded images of cardiomyocytes in ImageJ. Cumulative distributions of parameters were generated in MATLAB (MathWorks). A Kruskal–Wallis one-way analysis of variance test was used to compare the distributions. To compare individual parameters, 95% confidence intervals were determined by calculating the mean parameter value from 1000 bootstrap simulations. A test statistic was defined by calculating the difference in fit parameters between bootstrap simulations of the WT and mutant data, and the p-value was determined from the fraction of test statistic simulations that did not include the null hypothesis (i.e., that the parameters for the WT are the same as the mutant).

### Device fabrication for traction force microscopy

Hydrogels for traction force microscopy were generated in 35 mm dishes according to the protocol of Ribeiro et al., (21). The 14 mm glass-bottom of the 35 mm tissue culture dishes (MatTek) were silanized with 0.4% 3-trimethoxylyl-propylmethacrylate (Sigma) for 1 hour and then washed with water and dried with nitrogen gas. To make 10 kPa polyacrylamide hydrogels, a prepolymer solution containing 0.1% bisacrylamide was used (99). 0.2 μm diameter fluorescent beads were added to the prepolymer solution (FSSY002, Bangs Beads) as markers for the traction force.

Patterned coverslips were prepared using PDMS stamps and microcontact printing. 12mm round coverslips were cleaned by plasma cleaning under vacuum for 5 minutes. 1:10 dilution of Geltrex was prepared in DMEM-F12. 40 μL of Geltrex was added to the surface of the PDMS stamp and allowed to incubate for 20 minutes. Then the stamp was dried with nitrogen gas and placed on the 12 mm round coverglass for 2-3 minutes under a 50 g or 100 g weight, transferring the pattern to the coverglass. Unpatterned coverslips were prepared by coating the entire surface of a 12 mm coverglass with Geltrex.

To assemble the hydrogels, 15 μL of prepolymer solution was added to the silanized glass tissue culture dishes and then the 12 mm Geltrex-patterned coverglass was placed pattern side down. Hydrogels were allowed to polymerize for 15 minutes and then PBS was added for at least 1 hour to hydrate the hydrogels. Hydrogels could also be stored in PBS at 4°C until cell seeding. Before cell seeding, the 12 mm coverglass was removed using forceps.

### Traction force microscopy

All traction force microscopy measurements were performed on patterned polyacrylamide hydrogels with a stiffness of 10 kPa with cardiomyocytes that were at least 30 days post-differentiation. Cardiomyocytes were singularized using 0.25% trypsin and then resuspended in RPMI-20 with 10 μM Y-27632. PBS from the hydrogels was aspirated and 100 μL of media containing 50,000 cells was added to the hydrogel surface. After 45 minutes, 2 mL of RPMI-20 with 10 μM Y-27632 was added to the well. Cells were placed in the incubator for 48 hours, and then the media was changed to RPMI/B27+I and changed every 3 days until imaging.

On the day of imaging, fresh RPMI/B27+I was added 1 hour before imaging. Single, spontaneously-beating cells were imaged in an environmentally controlled chamber (Tokai Hit) on a Nikon spinning disk confocal microscope equipped with a Yokagawa CSU-X1 variable speed Nipkow spinning disk scan head and Andor Zyla 4.2 Megapixel sCMOS camera (Washington University Center for Cellular Imaging). Cells were imaged over an 800×400 pixel region of interest using a 40x objective. 20 seconds of data were collected at 33 frames per second. Both bright-field and fluorescence images were recorded.

### Analysis of traction force microscopy data

Traction force movies were analyzed by generating displacement maps of the fluorescent beads using a MATLAB program developed by the Pruitt lab (56). The program enables the measurement of several parameters, including the force and power output of single cells. The average rates of force development and relaxation for each cell were calculated by fitting the time courses of the force developed with single exponential functions. Cumulative distributions of parameters were generated in MATLAB. A Kruskal–Wallis one-way analysis of variance test was used to compare the distributions. To compare individual parameters, 95% confidence intervals were determined by calculating the mean parameter value from 1000 bootstrapped simulations. A test statistic was defined by calculating the difference in fit parameters between bootstrapping simulations of the WT and mutant data, and the p-value was determined from the fraction of test statistic simulations that did not include the null hypothesis (i.e., that the parameters for the WT are the same as the mutant).

### Hydrogel fabrication for immunofluorescence microscopy

Hydrogels for immunofluorescence microscopy were generated on 12 mm glass coverslips according to the protocol of Ribeiro et al., (21). 12mm glass coverslips were silanized with 0.4% 3-trimethoxylyl-propylmethacrylate (Sigma) for 1 hour and then washed with water and dried with nitrogen gas. To make 10 kPa polyacrylamide hydrogels, a prepolymer solution containing 0.1% bisacrylamide was used (99).

Patterned coverslips were prepared using PDMS stamps and microcontact printing. 12 mm round coverslips were cleaned by plasma cleaning under vacuum for 5 minutes. 1:10 dilution of Geltrex was prepared in DMEM-F12. 40 μL of Geltrex was added to the surface of the PDMS stamp and allowed to incubate for 20 minutes. Then the stamp was dried with nitrogen gas and placed on the 12 mm round coverglass for 2-3 minutes under a 50 g or 100 g weight, transferring the pattern to the coverglass.

To assemble the hydrogels, 15 μL of prepolymer solution was added to the silanized 12 mm coverglass and then the 12 mm Geltrex patterned coverglass was placed pattern side down to create a sandwich between the two coverglasses. Hydrogels were allowed to polymerize for 15 minutes and the sandwiches were transferred into a 24-well tissue culture plate. PBS was added for at least 1 hour to hydrate the hydrogels. Hydrogels could also be stored in PBS at 4 °C until cell seeding. Before cell seeding, the top coverglass was removed using a razor blade, leaving behind the 12 mm coverglass with the patterned hydrogel. Cell seeding and immunofluorescence preparation and imaging were carried out in the same manner as described in the traction force microscopy and immunofluorescence sections above.

---

**Figure 1 – figure supplement 1:**
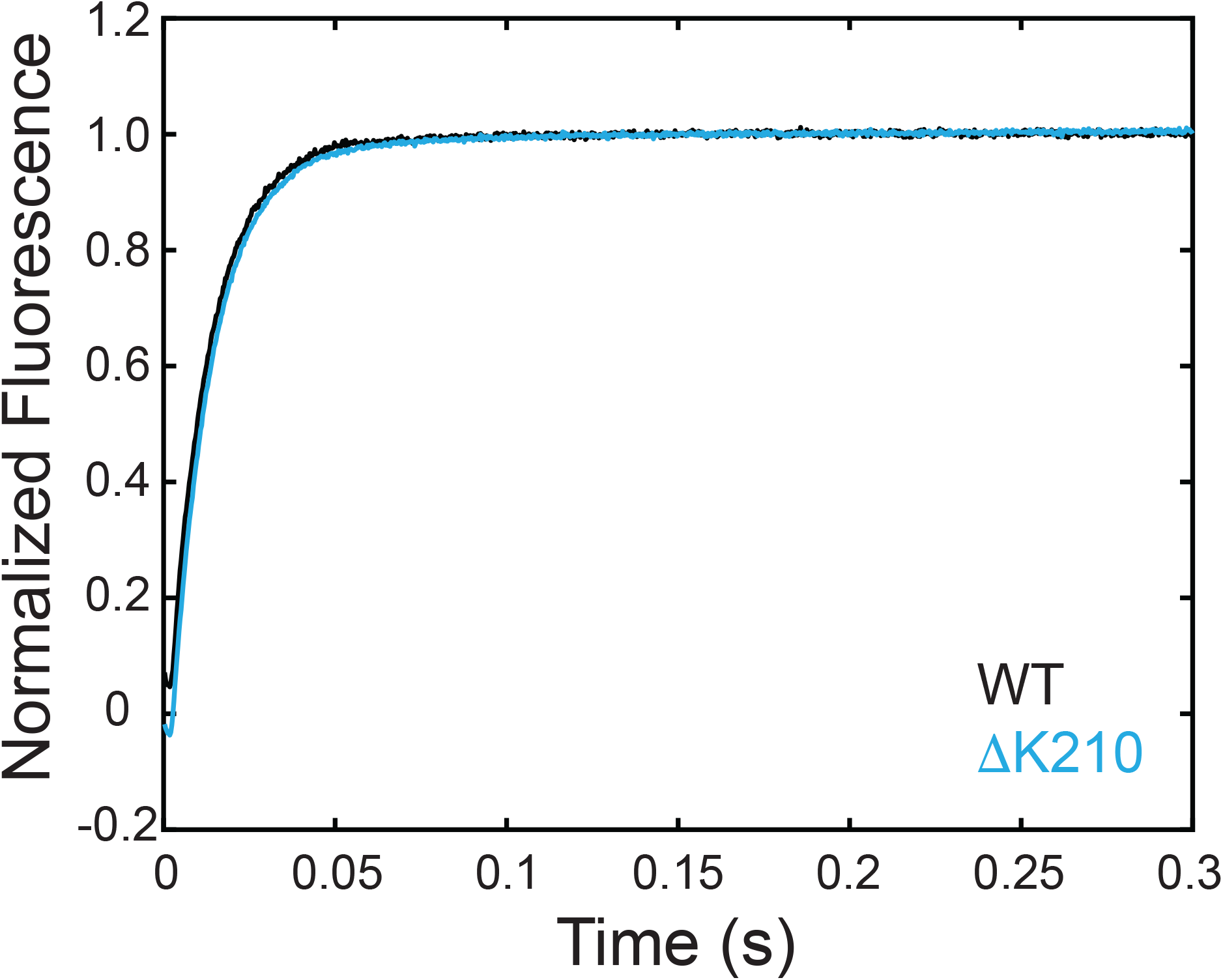
ADP release rates from myosin bound to regulated thin filaments for WT and ΔK210 at 20°C. The data is the average of at least three individual experiments. The ADP release rate in the presence of the mutation (72 s^-1^) is not significantly different from the WT (76 s^-1^, p=0.344).

**Figure 5 – figure supplement 1:**
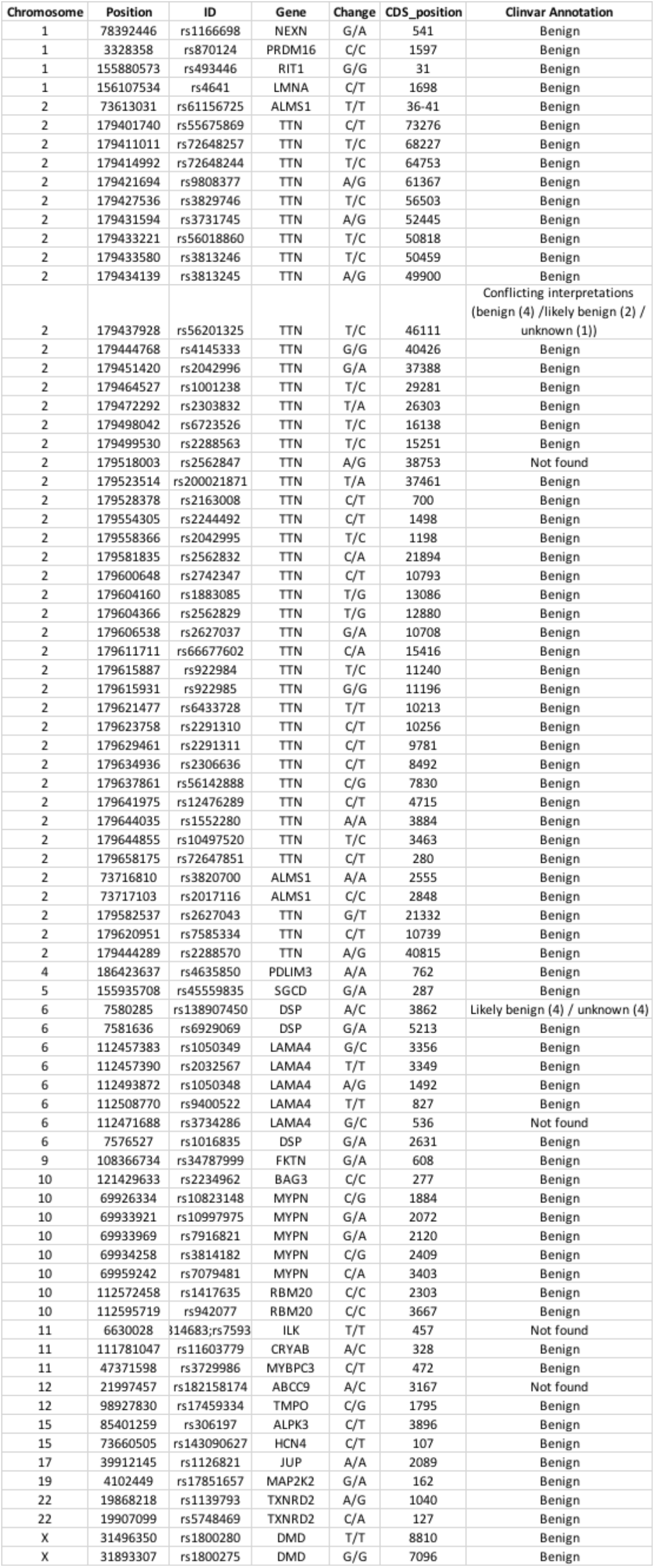
Variants identified from whole exome sequencing of BJFF cells. None of the identified variants in the BJFF line are classified as pathogenic or likely pathogenic in the ClinVar database.

**Figure 5 – figure supplement 2:**
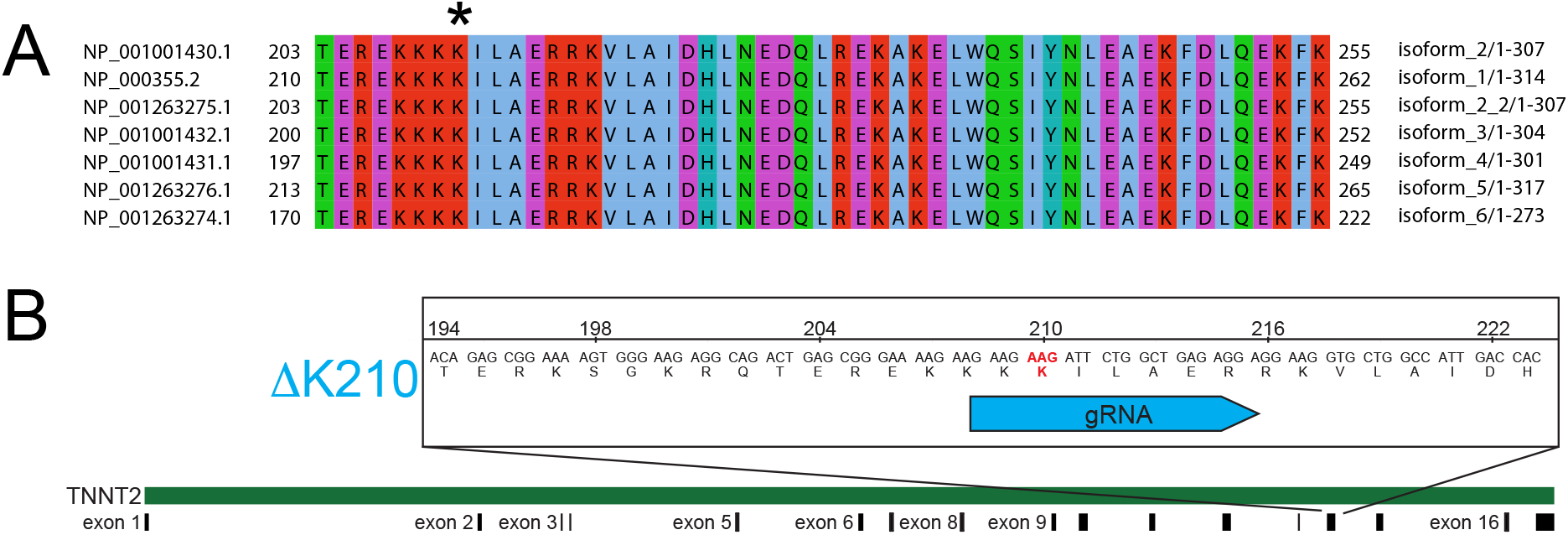
(A) Sequence alignment of all expressed human TNNT2 splice isoforms. K210 (*) is located in a conserved region of the protein. (B) CRISPR/Cas9 constructs for introducing the mutation into stem cells. This gRNA was chosen to minimize the risk of potential off-target effects. The AAG deletion was generated using a separate oligo for homology-directed repair, with an efficiency of ~10%. Two homozygous clones positive for the mutation were validated using Next Generation Sequencing.

**Figure 5 – figure supplement 3:**
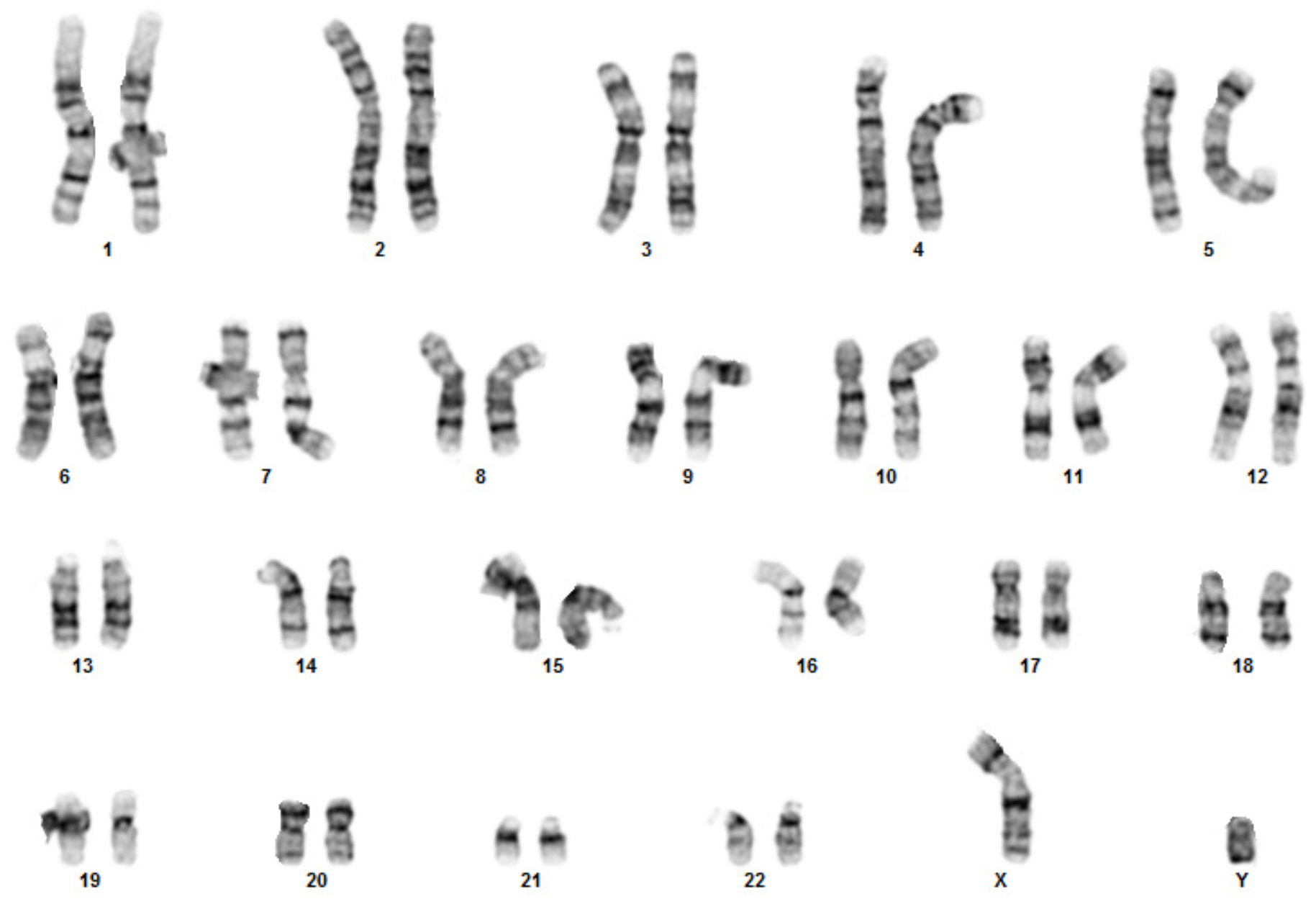
Karyotyping of gene edited cells. Cytogenetic analysis was performed on 20 G-banded metaphase cells (Cell Line Genetics) and these cells had normal karyotypes.

**Figure 5 – figure supplement 4:**
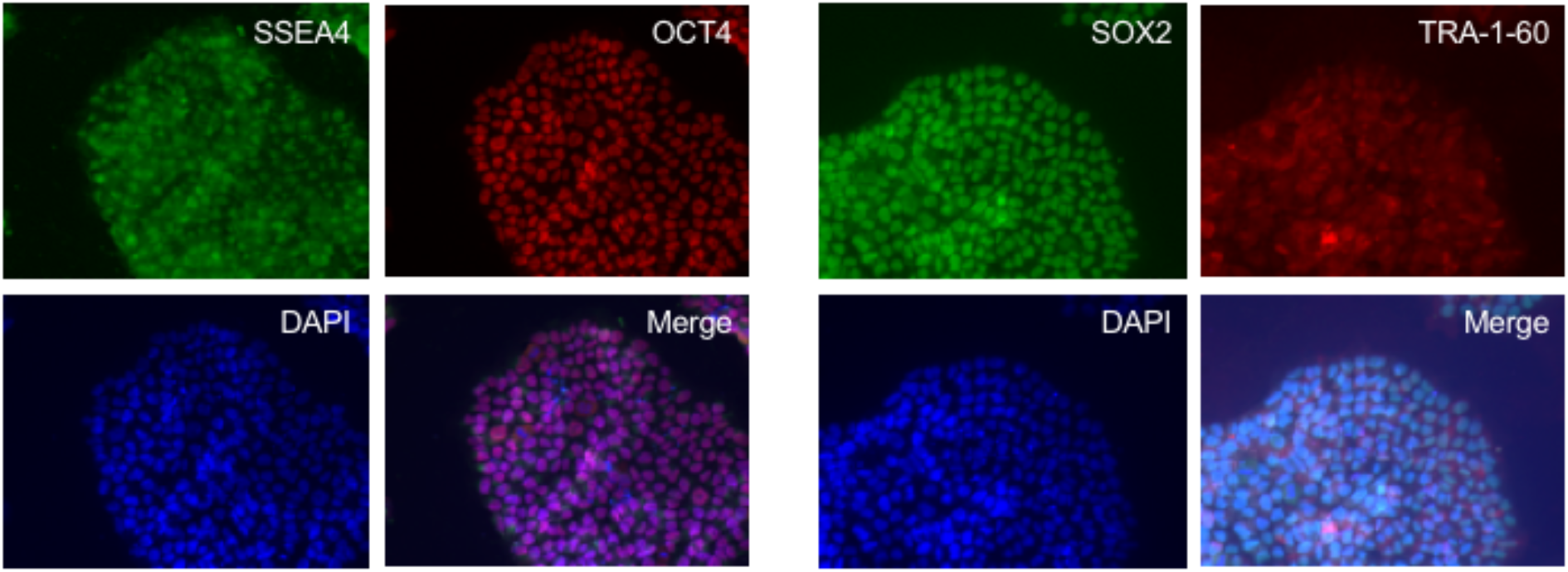
Staining of gene-edited cells with markers of pluripotency demonstrates that stem cells retain pluripotency after gene editing.

**Figure 5 – figure supplement 5:**
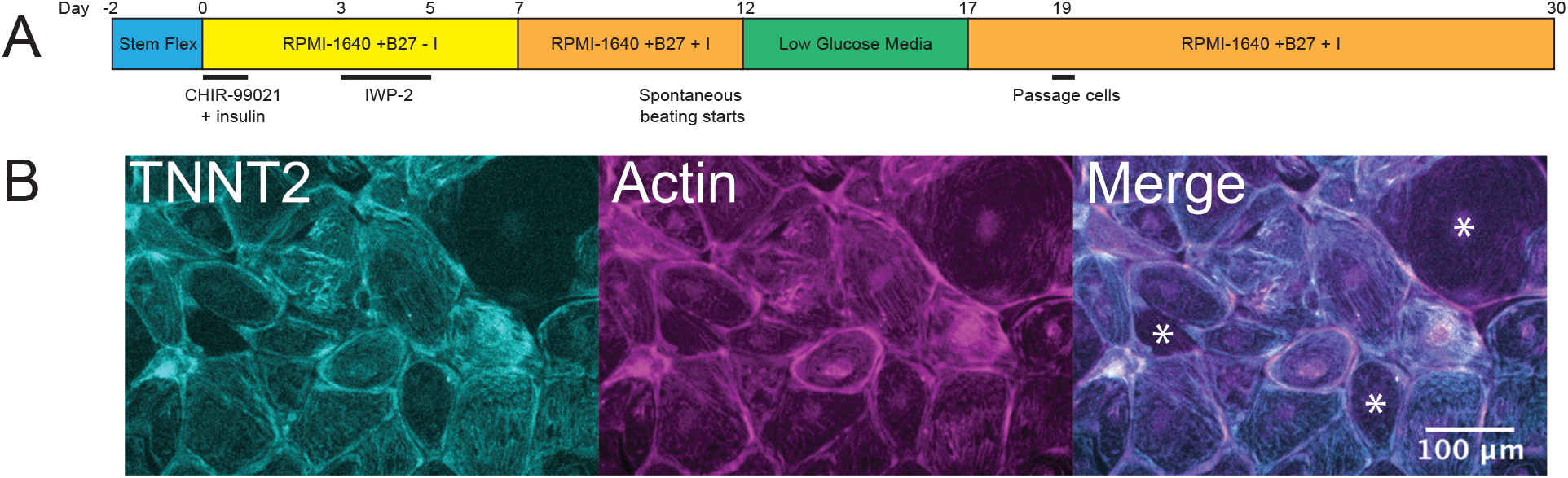
(A) Differentiation protocol used. I indicates insulin. (B) Cardiomyocyte purity was assessed by immunofluorescence. Cells were stained 30 days after differentiation to assess cardiomyocyte purity. Cells were stained using an antibody for cardiac troponin-T (TNNT2, cyan) and phalloidin which labels actin and acts as a marker of all cells (magenta). Cells were imaged using epifluorescence illumination. Non-cardiomyocytes contained actin, but not TNNT2 (denoted with *). 711 cells were observed, 652 of which were cardiomyocytes. This gives a cardiomyocyte differentiation efficiency of 92%.

